# Programmable bioprinting of tumor microenvironment arrays reveals laminin-dependent drug sensitivity

**DOI:** 10.64898/2026.08.21.745937

**Authors:** Marie Moulin, Edina Sehic, Adam Engberg, Christina Stelzl, Fredrik Holmberg, Tomas Bohn Pessatti, Benjamin Schmuck, Anna Rising, Johan Kreuger, Paul O’Callaghan

## Abstract

We present an active mixing toolhead for extrusion bioprinting. The tool enables the programmable fabrication of tumor microenvironment gradient arrays, through controlled deposition of mixed hydrogel precursor formulations into 384-well plates, pre-seeded with tumor cells. It operates on an open-source bioprinter and can actively mix arbitrary ratios of two hydrogel precursors prior to extrusion. These concentration gradient arrays are compatible with quantitative image analysis of cell viability and morphological responses to hydrogels conditioned with drug or extracellular matrix (ECM) proteins. The tools capacity to mix and print hydrogel precursor gradients was demonstrated using alginate and highly concentrated mCherry-conjugated mini-spidroin solutions. Hydrogel precursor stocks contained fluorescent reporters to facilitate quantifications of mixing efficiency, and as proxies for drug and ECM protein concentrations. The tool was applied to generate hydrogel-based gradients of the apoptosis-inducer staurosporine, from which concentration-dependent MDA-MB-231 breast cancer cell death responses were quantified. Gradient arrays of the ECM protein laminin-511, implicated in breast cancer tumorigenesis, were generated and revealed that increasing laminin-511 concentrations potentiated staurosporine-induced cell death. The study demonstrates the utility of this active mixing toolhead for producing hydrogel gradient arrays, and demonstrates the relevance of studying drug-responses in tumor microenvironment models that account for disease-specific ECM components.

## Introduction

Investigating how tissue microenvironments regulate cellular behavior is essential for the development of disease models and effective drug screening platforms^1,2^. *In vivo,* the extracellular matrix (ECM) varies greatly across tissues, and through interactions with cells plays central roles in development, tissue homeostasis and disease progression where its composition and mechanical properties are often remodeled^3,4^. During tumorigenesis, ECM remodeling correlates with poor prognosis, increased metastatic risks, and treatment resistance^5^.

Hydrogel-based models provide an *in vitro* system for evaluating how specific ECM compositions and mechanical properties of tissue microenvironments influence cell responses and drug sensitivity^1,2,6–10^. Alginate-based hydrogel precursors are widely used in bioprinting due to their simplicity and biocompatibility, providing a versatile matrix that can be readily conditioned with different molecules of interest. Chemically or genetically engineered biomaterials enable the design of hydrogel microenvironments with specific cell-adhesion motifs, tunable mechanical properties, and controlled spatial and temporal regulation of drug release or binding site exposure^11–14^. Spider silk proteins (spidroins), including engineered mini-spidroin variants, constitute an emerging class of advanced biomaterials that form cytocompatible hydrogels with controllable stiffness and can be co-expressed with diverse fusion proteins^15–20^.

To systematically and efficiently explore such complex microenvironments requires the generation of sequential iterations of hydrogel mixtures. Various methods have been developed to produce large compositional arrays of biomaterials^21–28^, which enable the investigation of cellular responses across a broad range of material formulations. Compositional hydrogel gradients can be generated through controlled mixing of multiple precursor solutions. This can be achieved with passive mixing systems, including microfluidic platforms where channel geometries promote molecular diffusion between laminar flow streams and in some cases induce chaotic advection^24,26,28–33^. In contrast, active mixing systems can facilitate mixing of hydrogel precursors without the need for continual flow. For example, a rotating impeller can be used to control the speed and duration of mixing, which can be particularly advantageous for high viscosity hydrogel precursors^34–36^. These approaches can generate continuous gradients in a variety of formats, including microarrays, hydrogel slabs, filaments and 3D constructs. Spatially resolved screens within such gradients can thereby attribute cell responses to specific properties of the hydrogel microenvironment. Such strategies have been applied to model healthy and tumor microenvironments, as well as study cell-biomaterial interactions^24,28,32^.

In this study, we present an active mixing toolhead that is integrated into the workflow of an open source extrusion bioprinter, which we previously described^37^. The tool consists of a mixing chamber supplied by two hydrogel precursor inlets that are respectively connected to independent programmable syringe pumps. These permit arbitrary mixing ratios to be defined and dynamically adjusted. Mixing is achieved via a centrally located rotating rod that is actuated using a stepper motor with adjustable rotation speed. The tool enables dynamic mixing of the two hydrogel precursor inputs, and spatially-controlled extrusion of discrete volumes. This functionality was applied to generate multi-well plate-based gradient arrays of hydrogels in which the concentration of an incorporated molecule of interest is incrementally varied. We characterize the tools capacity to form gradient arrays of alginate and mini-spidroin based hydrogels, using fluorescent reporters to quantitatively assess mixing efficiency. Hydrogel gradient arrays were then bioprinted directly onto MDA-MB-231 breast cancer cells pre-seeded in 384 well plates. Using a high-throughput image analysis strategy we quantitatively assess cell viability in gradient arrays of the pro-apoptotic agent staurosporine and calculate its half maximal effective concentration (EC_50_). To model the effects of a tumor microenvironment component that has been implicated in breast cancer progression we bioprinted gradient arrays of laminin 511 and determined that increasing concentrations of this ECM molecule correlated with increased MDA-MB-231 cell sensitivity to staurosporine.

The platform enables the generation of complex hydrogel gradient arrays, providing an *in vitro*, scalable approach for dissecting how variations in ECM composition influence cellular responses. By integrating active mixing with high-throughput screening, our system offers a versatile tool for modeling tumor and other disease-relevant microenvironments. This functionality aims to provide models that better account for how the extracellular niche impacts cell responses to specific therapeutic strategies.

## Results

### Mixing tool design and assembly

The mixing tool was designed to be operated using an open source bioprinter based on the E3D motion system and tool changer^37^. The mixing tool consists of three main 3D-printed parts, including the upper and lower housing, and the back mount. The back mount attaches the mixing tool through the upper housing to the E3D printer tool mount (Fig. 1A, and C Rear view). The stepper motor mounted on top of the upper housing (Fig. 1A and C) includes a drive shaft for coupling to the mixing shaft base. The mixing shaft base features threaded holes for three grub screws, which secure the attachment to the drive shaft (Fig. 1B). The lower housing comprises the mixing chamber, into which the mixing rod is inserted via an O-ring sealed opening (Fig. 1A and D). The bearing and the 3D-printed press lid ensure proper alignment of the mixing shaft base, thereby maintaining effective O-ring sealing of the mixing chamber around the mixing rod. The mixing chamber features two side inlets into which ferrule-capped tubing are inserted and O-ring sealed (Fig. 1A and D). The other end of each tube is connected using 3D-printed connectors to a syringe filled with hydrogel precursors. Each syringe is placed in a syringe pump tool, which has previously been described by Engberg et al.^37^. The syringe pump tools are programmable, and control the volume and ratio of hydrogel precursor extrusion into the mixing chamber. Centrally located at the bottom of the mixing chamber is the extrusion port, which is designed as a Luer lock receiver to fit an extrusion nozzle or needle (Fig. 1A, C and D). A photograph of the assembled mixing tool is presented in Fig. 1E. The design of the lower housing and the mixing shaft was refined during the study to optimize shaft alignment and to prevent leakage of hydrogel precursors at the O-ring seal. For example, the mixing tool base and rod was initially machined as a single solid piece, but the rod and shaft were subsequently produced as a two-part press-fit assembly, which improved perpendicular alignment, reduced runout, and permitted rod replacement if required.

**Fig. 1.**
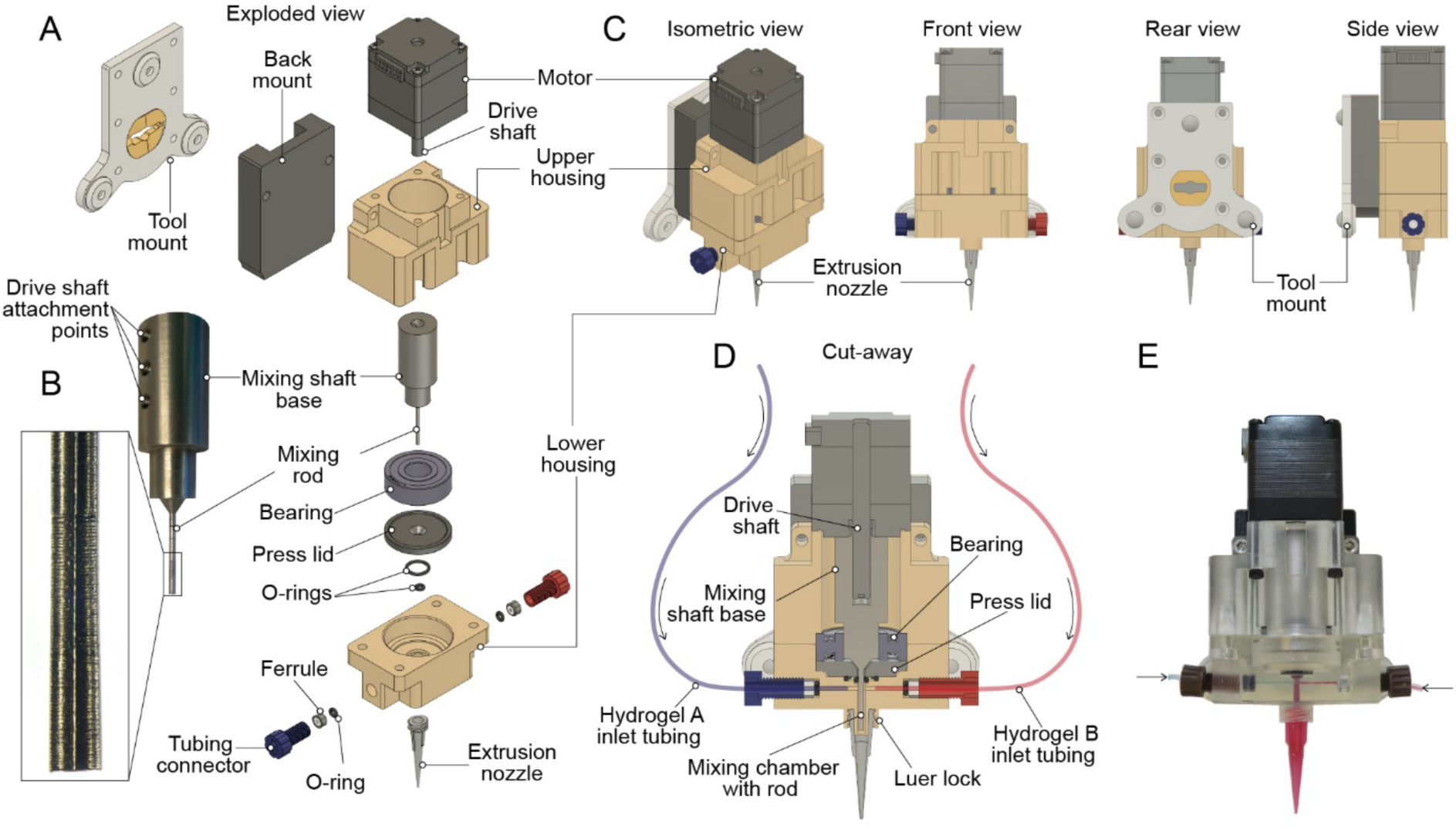
Mixing tool design and assembly. **A**. Exploded view of the CAD drawing of the mixing tool assembly. **B**. Photograph of the machined metal part that serves as the mixing shaft, which is attached to the drive shaft of the motor via the mixing shaft base, which has three threaded holes for attachment grub screws. In the zoomed in view, the profile of the ribbed surface of the machined mixing rod is highlighted. **C.** Four orientations of the CAD drawing of the assembled mixing tool. The isometric view details the position of the motor and rear tool mount plate. The front view highlights the tubing attachment points for the hydrogel precursor tubing inlets (blue and red), and the relative position of the extrusion nozzle. The rear view illustrates the lock point in the tool mount and the side view permits an appreciation of the tool’s relative height to depth ratio. **D.** Cut-away of the CAD drawing detailing the drive shaft inserted in the mixing shaft base; the mixing rod aligned in the mixing chamber with the bearing and press lid; the position of the O-rings; the assembled tube connectors within their respective inlets; and an attached extrusion nozzle. The tubing illustrates the flow of hydrogel precursor A (blue) and hydrogel precursor B (red) into the mixing chamber from their respective syringe pump tools. **E.** Photograph of the assembled mixing tool (3D printed with clear resin) filled with dyed blue alginate (hydrogel precursor A) and dyed red alginate (hydrogel precursor B).

### Mixing and extrusion of printed gradient arrays using fixed injection ratios of hydrogel precursors

The mixing tool is designed to mix two hydrogel precursors, referred to as hydrogel precursor A (Hy. A) and hydrogel precursor B (Hy. B), in the mixing chamber (Fig. 2A). Prior to printing, hydrogel precursors can be pre-conditioned with a specific molecule of interest (MOI), e.g. a pharmacological agent or an ECM molecule. The injected volumes and therefore ratio of hydrogel precursor A and hydrogel precursor B are independently determined by the two syringe pump extrusion tools (described previously in Engberg et al.^37^) that are programmed to simultaneously actuate the pistons of each hydrogel-containing syringe. As it is an open system, the total volume of hydrogel precursors A and B that is injected into the mixing chamber will simultaneously displace an equal volume of mixed hydrogel precursors, which will be extruded through the nozzle. The available volume in the mixing chamber, with the mixing rod accounted for, was calculated from the CAD file to be approximately 40 µl. The chamber is continuous with the downstream attached nozzle, and we typically used a 25 G nozzle with an available volume of approximately 160 μl when connected via the Luer lock. Therefore, the total volume in which hydrogel precursor mixing can occur in the toolhead was estimated to be 200 μl (Fig. 2A). In Fig. 2B, we predict the gradient array that would be generated using a 50:50 fixed injection ratio of hydrogel precursors A (depicted in blue) and B (depicted in red). The mixing chamber and nozzle are prefilled with 200 µl of hydrogel precursor A. As outlined in Fig. 2B, the first extrusion (E1) occurs simultaneously with the injection of 10 µl of hydrogel precursor A and 10 µl of hydrogel precursor B into the mixing chamber; consequently, E1 will be 20 µl in volume and consist of 100% hydrogel precursor A. Assuming the injected hydrogel precursors A and B are homogenously mixed as part of the total 200 µl volume, this will result in a dilution of hydrogel precursor B such that it constitutes 5% of the mixture. This defines the composition of extrusion two (E2), which will be displaced with the next 50:50 injection of hydrogel A and B. With this fixed injection ratio program, we can produce an extrusion series for a hydrogel precursor mixture in which the hydrogel precursor A component is sequentially diluted, while hydrogel precursor B is sequentially concentrated. The expected profile for forty extrusions of this fixed 50:50 mixing ratio are presented in Fig. 2C. The resulting plot fits to a one-phase exponential model, and outlines how the percentage of the MOI-containing hydrogel precursor (hydrogel B in the example) would initially increase with a near linear profile, but then gradually level off as it approaches the intended final concentration of 50% (49.87% in E40, Fig. 2C).

**Fig. 2.**
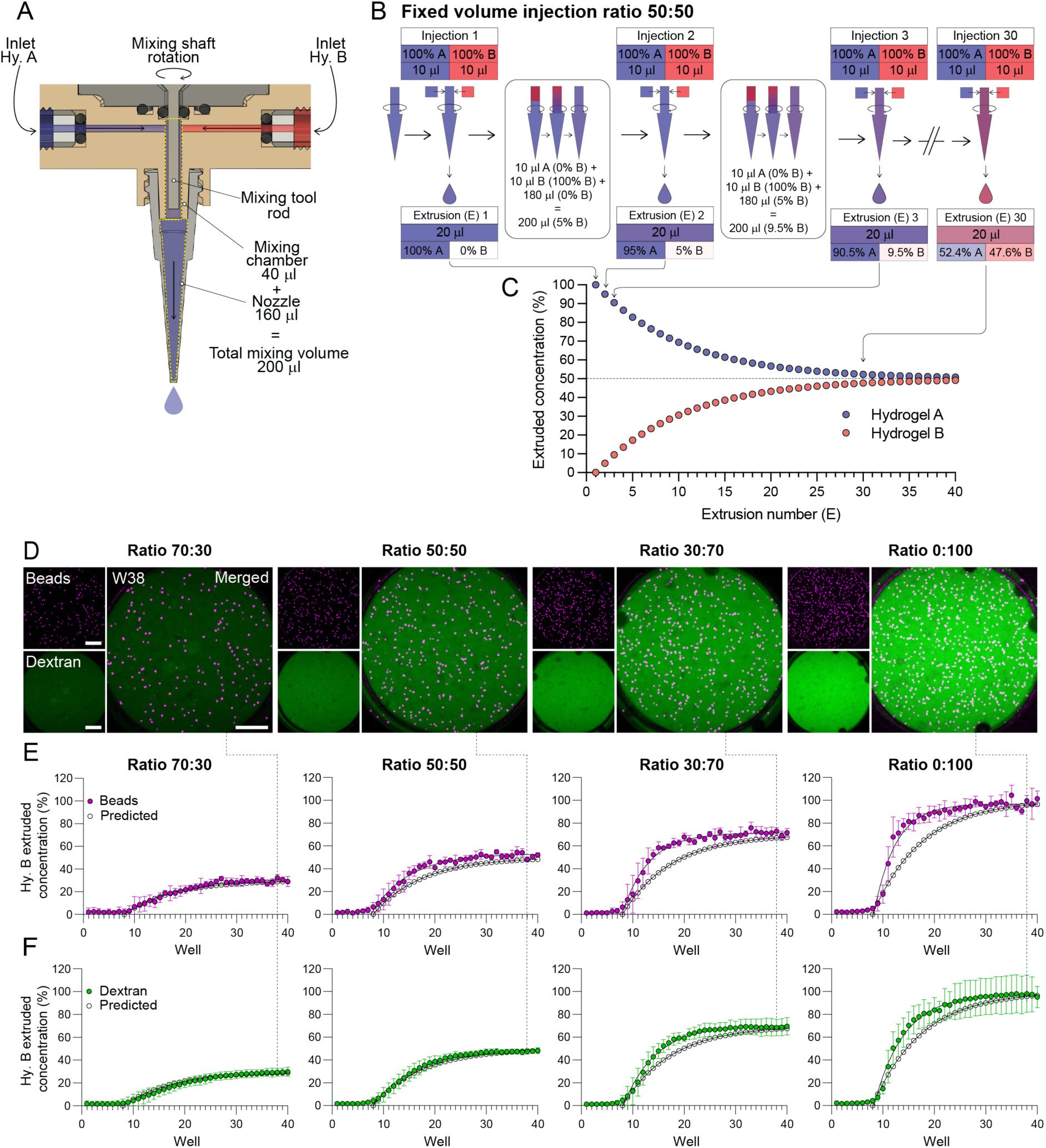
Gradient generated by mixing tool prints with fixed injection ratios. **A.** Cutaway view of the design of the mixing chamber, outlining the total 200 µl mixing volume as defined by the mixing chamber and attached extrusion nozzle. Hydrogel precursor A (Hy. A) is depicted in blue and hydrogel precursor B (Hy. B) is depicted in red. **B.** Graphical representation of the injection, mixing and extrusion protocol and theoretical outcomes when mixing hydrogel precursor A (blue) and hydrogel precursor B (red) with a 50: 50 injection ratio. The mixing tool is prefilled with hydrogel precursor A. The syringe pumps are programmed to inject 40 sequential injections of 10 μl of hydrogel precursor A and 10 μl of hydrogel precursor B into the mixing chamber, each resulting in the simultaneous extrusion (E) of 20 μl. Extrusion numbers 1, 2, 3 and 30 are illustrated in the example. Their expected concentrations are mapped to their relative positions on the theoretical plot in **C.** Theoretical plot outlining the extruded hydrogel precursor mixtures with their respective concentrations of hydrogel precursor A and hydrogel precursor B for 40 consecutive extrusions for a 50:50 fixed injection ratio. **D.** Images of the CountBright fluorescent microspheres (magenta) and dextran Alexa Fluor 488 (dextran-AF488; green) in extrusion well 38 for each of the indicated fixed injection ratios. **E-F.** Fixed injection ratio print profiles for Hy. A: Hy. B (70:30, 50:50, 30:70, 0:100) extruded into 40 wells. Hydrogel precursor B was pre-conditioned with dextran-AF488, and with CountBright fluorescent microspheres. (E) Relative numbers of fluorescent microspheres (beads) and (F) relative dextran-AF488 (dextran) intensity in each well expressed as percentages of the average bead number and fluorescence intensity in control wells with only hydrogel precursor B. Experimental data are plotted for all of the indicated fixed injection ratios overlaid with their predicted extrusion profiles represented with white dots (calculated as in Fig. 2B)). Scale bar in D: 500 μm. Datapoints in E and F represent the mean ± SD, derived from n = 4 independent printed arrays.

### Experimental evaluation of extruded hydrogel precursor concentration gradients

Next, we experimentally characterized the extrusion profiles for a selection of gradient arrays that were generated using different fixed injection ratios for hydrogel precursors A and B, and compared them with their equivalent predicted profiles. To track the concentration of hydrogel precursor B in each extrusion, we pre-conditioned it with two reporters: a suspension of fluorescent microspheres (beads), and 10 kDa dextran conjugated to the fluorophore Alexa fluor 488 (dextran-AF488). Hydrogel precursor A served as the diluent hydrogel precursor and contained neither of the fluorescent reporters. The operation of the mixing tool on the E3D printer facilitates automatized displacement of the tool in the (x, y) direction and print platform displacement in the z direction. Therefore, each extrusion can be deposited into an individual well of a 384 well plate. Each fixed ratio print was programmed to produce a series of 20 μl extrusions into 40 wells. Confocal microscopy images from the wells were then acquired and analyzed to quantify bead numbers and dextran-AF488 fluorescence. Images of beads (magenta) and dextran-AF488 (green) fluorescence from a representative well (well 38) for each of the fixed ratio prints is presented in Fig. 2D. The quantified bead and dextran-AF488 concentrations for the 40 well extrusion series performed for each of the fixed injection ratio arrays of hydrogel precursor A and B are presented in Fig. 2E and F. Extruded concentrations of Hy. B were expressed as percentages of beads or dextran-AF488 fluorescence by normalizing fluorescence intensity and bead counts to control wells containing only hydrogel precursor B loaded with the fluorescent reporters (i.e. 100%). Unlike the predicted profiles, the extrusion profiles for the beads and dextran-AF488 fluorescence revealed an initial lag phase of approximately 7 wells during which no increase in the concentration of hydrogel precursor B (%) was detected. Potential explanations for this effect are discussed later. Overall, the subsequent fluorescence profile for the exponential and plateau phases of the extrusions correlated well with the predicted curves (Fig. 2E), suggesting that the tool is operating as expected and outlined in Fig. 2B and C. The extrusion profiles for the fluorescent bead counts exhibited more variability than the dextran-AF488 fluorescence (Fig. 2E and F). The smoother dextran-AF488 curves are likely due to the fact that the values represent average fluorescence from each extrusion well, while for the beads, each datapoint represents the number of individual beads detected. Noticeably, the fluorescence gradients that best fitted with the predicted curves were observed for prints using the lower ratios of hydrogel precursor B (i.e. 30% and 50% in Fig 2E and F). As the hydrogel precursor B component of the fixed ratio increased i.e. to 70% and 100%, the bead and dextran-AF488 fluorescence increased more steeply and plateaued earlier than predicted.

### Mixing and extrusion of printed gradient arrays using dynamic injection ratios of hydrogel precursors

To obtain a wide range of sequential changes in resulting hydrogel concentrations we generated a linear gradient array program, which was designed to incorporate the linear region from the profiles of each of the different fixed injection ratios. Fig. 3A illustrates how changing the injection ratios at fixed intervals during the extrusion program would be expected to produce an approximately linear gradient, with a range of 0-100% for hydrogel precursor B content. To further smoothen this gradient, a dynamic injection ratio strategy was designed such that after every four extrusions the Hy. A: Hy. B injection ratio would change to progressively increase the concentration of hydrogel precursor B in the extrusion (Fig. 3B). As before, the chamber was prefilled with 100% hydrogel precursor A, and the first injection ratio of 85:15 (17 µl:3 µl) and second to last injection ratio 15:85, were defined by the fact that 3 µl was the lowest hydrogel precursor volume that could be reliably injected with the syringe pump tool. The same experimental procedure used to characterize the fixed-ratio injection prints in Fig. 2 was applied to evaluate the dynamic gradient prints. The printed arrays were imaged using a confocal microscope with an automated stage to quantify fluorescent microsphere counts and dextran-AF488 fluorescence in each extrusion well (Fig. 3C). The dextran-AF488 fluorescence in the gradient arrays was also analyzed using a fluorescence plate reader, demonstrating the compatibility of these printed arrays with a standard downstream analysis platform (Fig. 3C). The quantified fluorescence in each extrusion produced profiles that closely matched the predicted dynamic gradient, but again required the predicted profile to be shifted to account for a seven-well delay in the detected change of the extruded hydrogel B concentration.

**Fig. 3.**
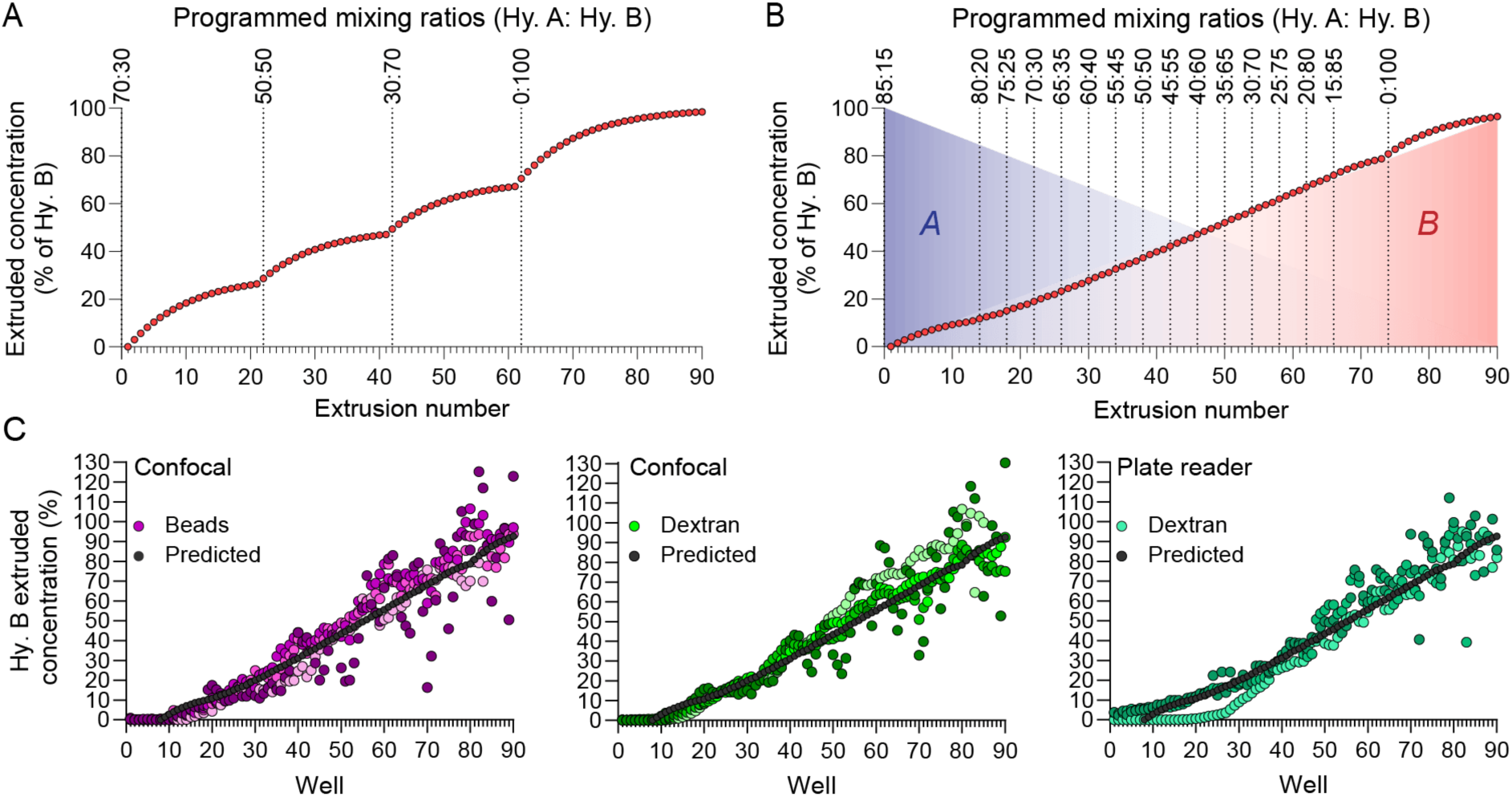
Linear gradients generated with dynamic injection ratios. **A.** Sequential alterations to the four fixed injection ratios of hydrogel precursor A (Hy. A): hydrogel precursor B (Hy. B), described in Fig. 2, are expected to produce a 0-100% range of extruded hydrogel B concentrations that approximate a linear gradient profile. **B.** Expected extrusion profile for hydrogel precursor B concentrations when the programmed Hy. A: Hy. B ratio is incrementally adjusted during printing. Injection ratios are changed every four wells, except at the beginning and end of the print. **C.** Experimental extruded concentrations of hydrogel precursor B during a dynamic injection ratio print as described in B, tracked using CountBright fluorescent microspheres (beads) and dextran-AF488 (dextran). Datapoints in C, collected using confocal microscopy represent the fluorescence per well from *n* = 4 independent printed arrays, and from *n* = 3 independent printed arrays analyzed with the plate reader. Data from each independent replicate is presented with a different shade of each color. The fluorescent bead and dextran-AF488 profiles are overlaid with the predicted concentration profiles for the dynamic injection ratio program (B), represented by black dots.

### Dynamic gradient arrays printed with the mini-spidroin protein A_3_I-A

To assess the mixing tool’s ability to produce gradients with a class of protein-based hydrogel precursors we prepared arrays of mini-spidroin solutions. Spidroin-based biomaterials (fibers and hydrogels) have a wide range of biomedical applications, including as cell culture microenvironments^16,17,38,39^. Recombinantly produced mini-spidroins can be expressed as chimeric proteins conjugated with enzymes or fluorescent reporters to produce different types of functionalized hydrogels^18,40^. Controlling the concentrations of such hydrogels also permits the tuning of their mechanical properties^16^; therefore, we next investigated the mixing toolheads capacity to generate gradient arrays of the mini-spidroin (A_3_I)₃-A_14_, hereafter referred to as (A_3_I-A)^41^. This construct is an engineered variant of the NT2RepCT mini-spidroin^15^ containing two poly-alanine blocks, with isoleucine substitutions introduced at every fourth position in the first block. A_3_I-A was combined in several configurations for gradient prints with the dynamic injection ratios mixing program. High concentration (200 mg/ml) A_3_I-A solutions were injected both as hydrogel precursor A and as hydrogel precursor B, with the latter stock first pre-conditioned with dextran-AF555, acting as a fluorescent reporter. Quantitative image analysis of the dextran-AF555 in the 90 extrusion wells revealed an increasing concentration gradient profile with a slope that was similar as that expected for a gradient array programmed for dynamic mixing ratios (Fig. 4A). Next, we tested the tools capacity to perform multi-material mixing, by injecting alginate (5% w/v), and combining it with a fusion protein consisting of A_3_I-A and mCherry^18^ (A_3_I-A-mCherry; 296 mg/ml). The A_3_I-A-mCherry color gradient was macroscopically visible in the printed extrusion array, and increasing mCherry fluorescence was detectable across the gradient by confocal imaging (Fig. 4B). Quantitative image analysis of mCherry fluorescence for each extrusion across the gradient revealed a linear increase, but with a steeper slope than that predicted. Consequently, the maximum (296 mg/ml) intended A_3_I-A-mCherry concentration was detected by well 50 instead of well 90, and from well 60 onwards, a number of extrusions failed to be deposited into their respective wells (Fig. 4B and C). A heatmap representation of the mCherry fluorescence per well across the printed array confirmed the generation of an incrementally increasing concentration gradient; however, a non-homogenous distribution of the A_3_I-A-mCherry component was observed within each well (Fig. 4D). Finally, we produced a decreasing concentration gradient array consisting of mixtures of the highly concentrated A_3_I-A-mCherry solution (256 mg/ml) with an unconjugated lower concentration A_3_I-A solution (83 mg/ml) (Fig. 4E). Quantitative image analysis of A_3_I-A-mCherry fluorescence across the 90 well extrusion array revealed the intended decreasing concentration gradient, but again deviated in its profile from the expected linear regression. Specifically, there was a delay in the initial decrease of A_3_I-A-mCherry concentrations, followed by a steeper decline in the gradient from approximately extrusion 70 onwards (Fig. 4E). Similar to Fig. 4D, the heatmap array corresponding to the decreasing A_3_I-A–mCherry fluorescence also revealed variability in its intrawell distribution (Fig. 4F).

**Fig. 4.**
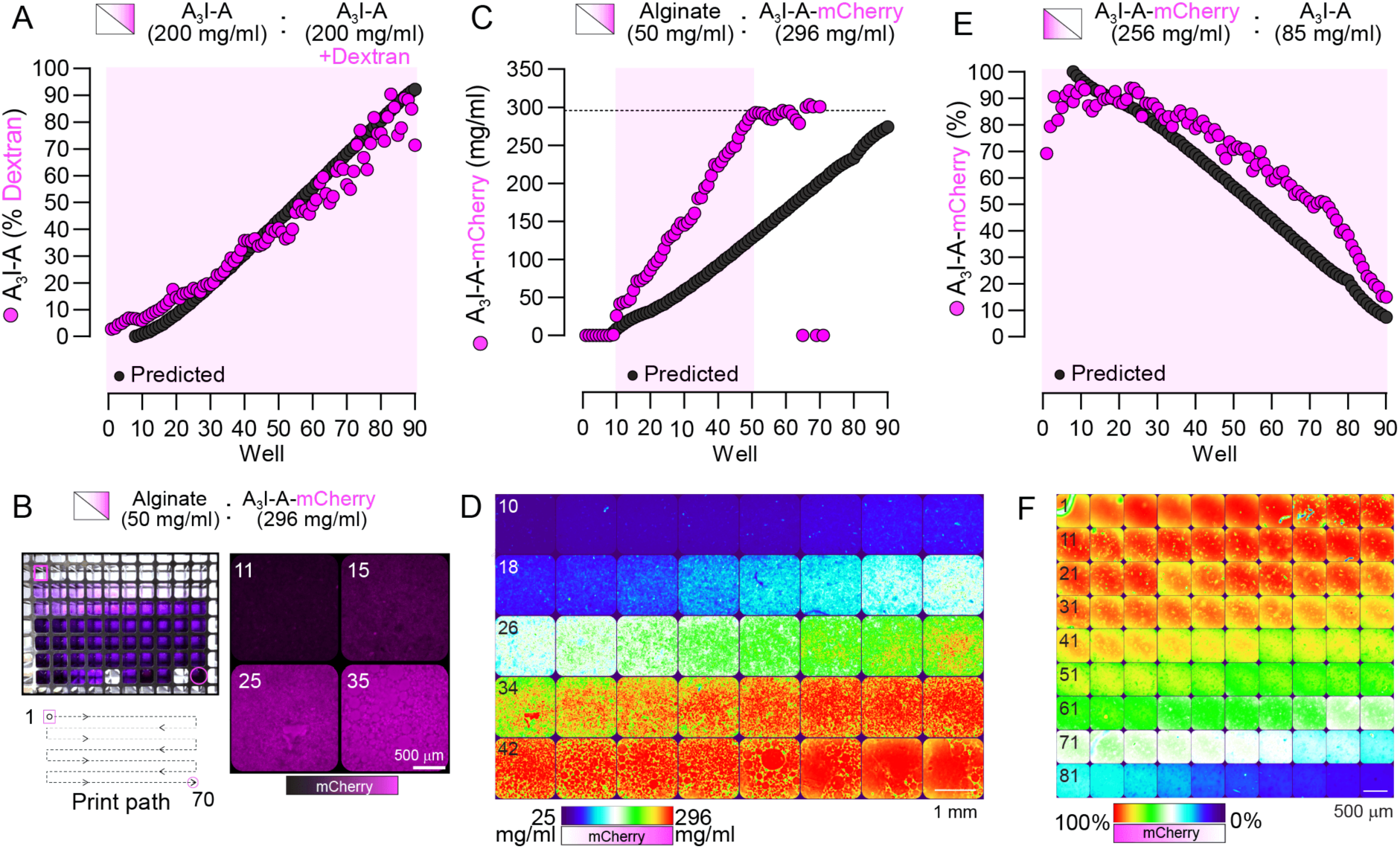
Dynamic gradient printing of A_3_I-A mini-spidroin solution arrays. **A.** Quantification of dextran fluorescence across sequential extrusions for A_3_I-A (200 mg/ml) mixed with a separate stock of A_3_I-A (200 mg/ml) containing dextran Alexa Fluor 555 (AF555) (10 kDa; 5 µM). Imaging was performed using a plate reader. **B.** Images of a bioprinted array of alginate (50 mg/ml) mixed with A_3_I-A–mCherry (296 mg/ml) and extruded into a 384-well plate. The serpentine print path is indicated. Right: confocal images of A_3_I-A-mCherry fluorescence from selected extrusions (11, 15, 25, 35). **C.** Quantification of A_3_I-A–mCherry concentrations across extrusions from the array in B. Concentrations were calculated from mCherry fluorescence recorded in extrusions of A_3_I-A-mCherry (296 mg/ml) only. Imaging was performed using confocal microscopy. **D.** Heatmap of mCherry fluorescence across the linear gradient region (extrusions 10–49; shaded magenta in C). **E.** Quantification of relative mCherry signal (%) across extrusions for A_3_I-A–mCherry (256 mg/ml) mixed with A_3_I-A (85 mg/ml) using a reversed version of the dynamic gradient print program. Imaging was performed using confocal microscopy. **F.** Heatmap of A_3_I-A-mCherry fluorescence across the entire printed array (extrusions 1–90; shaded magenta in E). In (A, C, E), black points indicate the predicted fluorescence or spidroin concentration for the dynamic injection ratio program.

### Screening MDA-MB-231 cell responses to increasing staurosporine concentrations in alginate hydrogel arrays

To demonstrate the platforms capacity to create defined drug gradients for screening dose-responses, we applied the dynamic injection ratio approach to print a linear gradient of the apoptosis-inducing drug staurosporine onto cultures of MDA-MB-231 breast cancer cells. Hydrogel precursor B was supplemented with 20 µM staurosporine, a protein kinase inhibitor known to induce apoptosis^42^, and dextran-AF488, which was used as a fluorescent proxy for calculating staurosporine concentrations extruded in each well (Fig. 5A). One day prior to printing, suspensions of the MDA-MB-231 breast cancer cell line were seeded into 384-well plates. A dynamic gradient print of increasing staurosporine concentrations in alginate was printed onto cells as 20 µL extrusions. Following gradient array printing, the hydrogels were crosslinked with CaCl₂ and maintained under standard culture conditions. Following 24 h of staurosporine exposure, the cells were subjected to live/dead staining with the Calcein AM (live) and propidium iodide (PI; dead) dyes. Calcein and PI fluorescence was imaged by confocal microscopy, and the percentage of dead cells relative to the total number of cells per well was quantified with CellProfiler software. The percentage of dead cells was plotted as a function of the staurosporine concentration in alginate hydrogels, which resulted in a sigmoidal dose–response curve, from which the EC_50_(death) of staurosporine-induced MDA-MB-231 cell death was determined as 1.01 ± 0.23 µM (Fig. 5B). In parallel, staurosporine-induced changes in cell morphology were quantified by measuring the aspect ratio (AR) of live cells and plotting this against staurosporine concentrations, with a calculated EC_50_(AR) of 0.86 ± 0.06 µM (Fig. 5C). Cells exposed to staurosporine concentrations below 0.86 µM (<EC_50_(AR)) and between 0.86 µM and 1.01 µM (>EC_50_(AR); <EC_50_(death)) are presented in Fig. 5D, where rounded morphologies are apparent in calcein-AM positive cells, prior to cells dying and becoming permeable to PI (Fig. 5D). This demonstrates the platforms utility for monitoring multiple cell responses in hydrogel gradient arrays, and correlating them directly with a specific drug concentration.

**Fig. 5.**
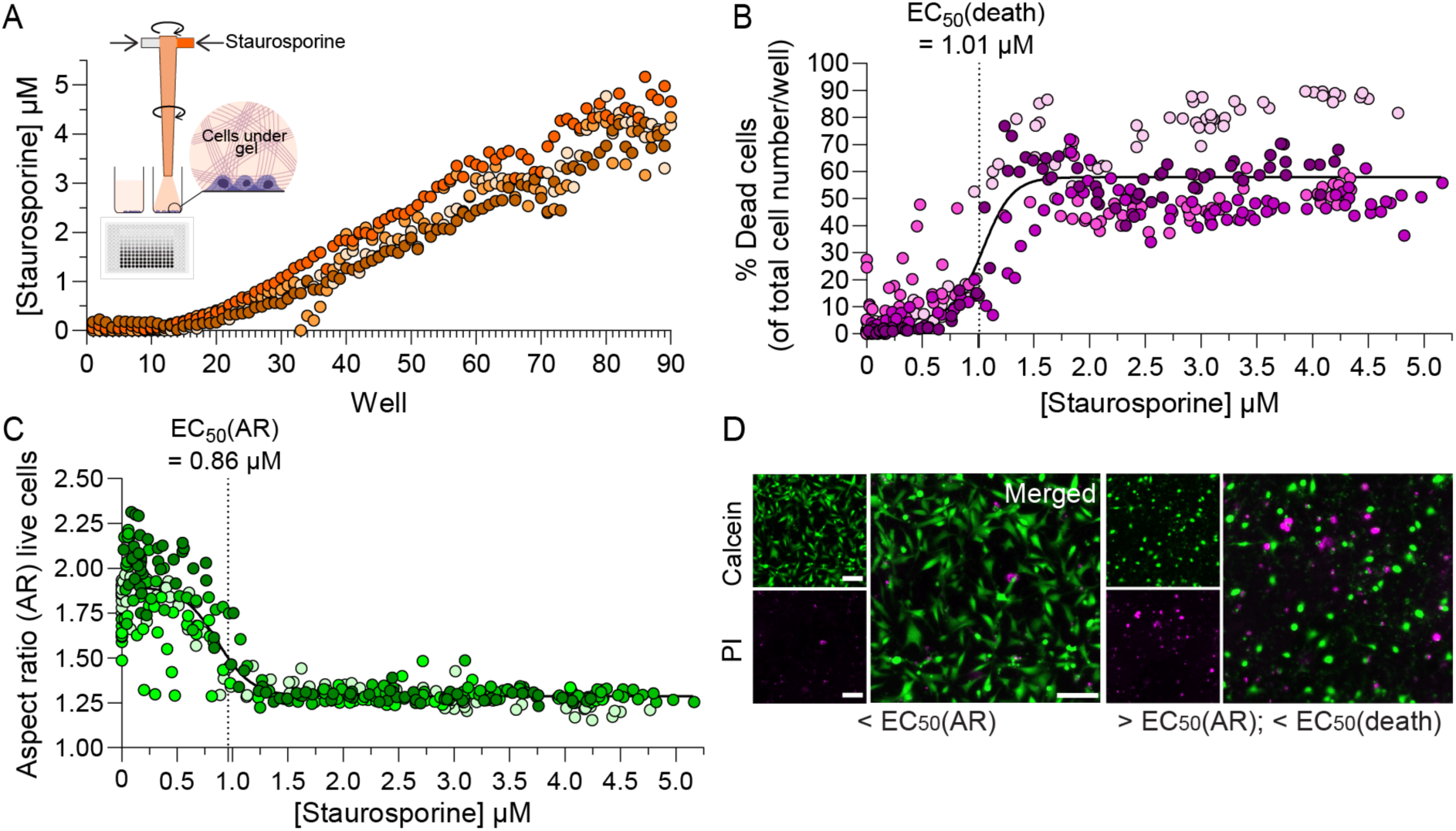
Screening MDA-MB-231 responses to a staurosporine gradient of concentration in alginate. **A.** Dynamic injection ratio gradient print of alginate (hydrogel precursor A) and staurosporine-conditioned alginate (hydrogel precursor B) extruded onto pre-seeded cultures of MDA-MB-231 cells in a 384 well plate. Staurosporine concentrations per well were tracked by confocal imaging and quantitative analysis of dextran-AF488 intensity, using dextran-AF488 intensity in hydrogel B only as the reference for maximal fluorescence intensity. The dynamic gradient of staurosporine in alginate is printed on MDA-MB-231 cells attached to the bottom of the wells. **B.** Quantification of staurosporine-induced death in MDA-MB-231 cells. Propidium iodide (PI)-positive (dead) cells were expressed as a percentage (%) of total cell numbers per well, as defined by the sum of Calcein AM- and PI-positive cells. Color-coded datapoints represent four independent biological replicates. An average half effective maximal concentration (EC_50_(death)) for staurosporine-induced MDA-MB-231 death was calculated to be 1.01 µM ± 0.23 µM (mean ± SD). **C.** Quantification of the change in aspect ratio (AR) of Calcein AM-positive MDA-MB-231 cells in each well of the staurosporine gradient. The data show the average AR for all Calcein AM-positive cells per well. The average EC_50_(AR) for staurosporine-induced changes in MDA-MB-231 aspect ratio was calculated to be 0.86 ± 0.06 µM (mean ± SD). The datapoints in (C) and (D) were fitted with a nonlinear regression curve using the log(agonist) vs. response model. EC_50_ values were determined from the fitted curves for each of four independent experiments, and the mean EC_50_ indicated in (C) and (D) was subsequently calculated from these individual values. **D.** Calcein AM and PI-positive MDA-MB-231 cells imaged with confocal microscopy from wells in the gradient array containing staurosporine concentrations that were below the EC_50_(AR) EC_50(AR)_ or above the EC_50_(AR) and below the EC_50_(death). (Scale bar in D: 100 µm).

### Screening effects of increasing laminin 511 concentrations on staurosporine-induced MDA-MB-231 cell death

Changes in the ECM content and mechanical properties of the tumor microenvironment (TME) have been attributed roles in tumor progression and in altered responses to anti-cancer agents^5,43^. For example, tumor-derived laminin 511 has been experimentally implicated in the metastatic progression of breast cancer^44^. Therefore, we next applied the dynamic mixing program to determine if fine-tuning the laminin 511 content in the alginate microenvironments alters MDA-MB-231 cell sensitivity to staurosporine-induced death. A linear gradient of laminin 511 in alginate was printed into 384-well plates pre-seeded with MDA-MB-231 cells. Laminin 511 concentrations were extrapolated from the fluorescence intensity of dextran-AF488, which was added to the stock hydrogel precursor conditioned with laminin 511 (Fig. 6A). Cells were exposed (24 h) to 1 µM staurosporine, which approximately corresponded to the EC_50_(death) determined in Fig. 5 for staurosporine-induced death in alginate only. The percentage of PI-positive dead cells were plotted against laminin 511 concentrations per well (Fig. 6B). Increases in the percentage of cell death induced by 1 µM staurosporine correlated with increasing laminin concentrations, with a significant positive slope (Fig. 6B, m_1_), while cells cultured below the laminin gradient arrays without staurosporine revealed low levels of PI-positive cells and a non-significant slope (Fig. 6B, m_2_). Changes in cell morphology were analyzed by plotting the aspect ratio of Calcein AM-positive (live) cells against laminin concentrations. In the absence of staurosporine, cells exhibited aspect ratio values > 1.6, indicative of an elongated morphology, though there was a slight, but significant decrease in the slope of the aspect ratio data along the laminin gradient (Fig. 6C, m_3_). Cells in the laminin-gradient treated with staurosporine revealed a lower aspect ratio, consistent with rounded morphology, which inversely correlated with laminin concentrations (Fig. 6C, m_4_). The percentages of staurosporine-induced MDA-MB-231 cell death were binned into three distinct laminin concentration ranges (0–1 µg/mL, 1–2 µg/mL, and 2–3.25 µg/mL), which similarly revealed that MDA-MB-231 cell sensitivity to staurosporine-induced death significantly increased with increasing laminin concentrations (Fig. 6D).

**Fig. 6.**
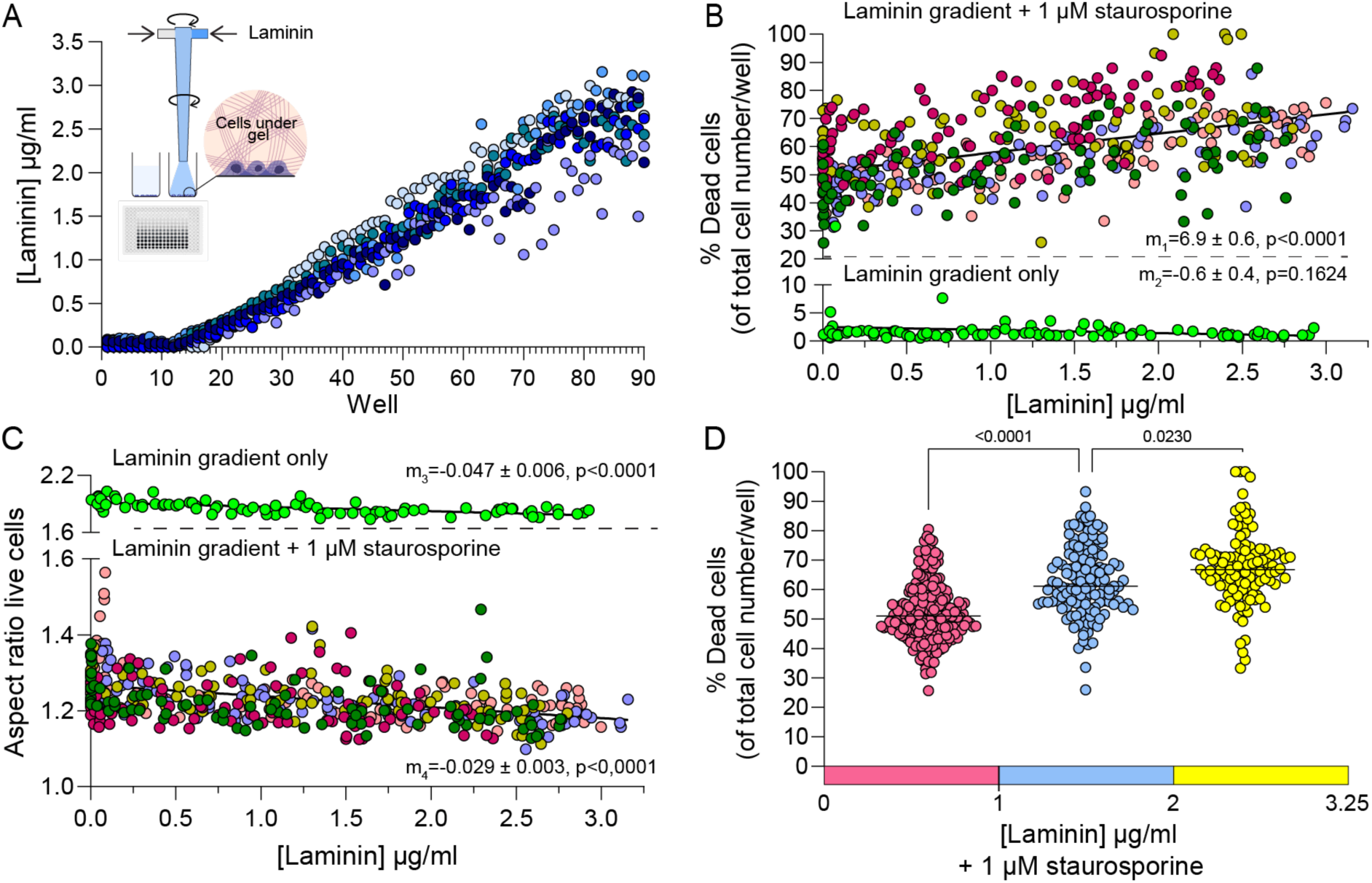
Screening MDA-MB-231 responses to a laminin 511 gradient of concentration in alginate with staurosporine treatment. **A.** Dynamic injection ratio print of alginate (hydrogel precursor A) and laminin 511 conditioned alginate (hydrogel precursor B) extruded onto pre-seeded cultures of MDA-MB-231 cells in a 384 well plate. Laminin 511 concentration per well is calculated from image analysis of dextran-AF488 fluorescence. **B.** Quantification of cell death in laminin 511 gradient arrays following exposure (24 h) to 1 µM of staurosporine, and an untreated control array. Color-coded plots represented data collected from four independent biological replicates. **C.** Quantification of the change in aspect ratio of Calcein AM-positive MDA-MB-231 cells in each well of the laminin gradient following exposure to 1 µM staurosporine, and an untreated control array. The data show the average aspect ratio for all Calcein AM-positive cells per well. For (B) and (C), the slopes (m) of the linear regressions, along with their standard deviation and corresponding p value, are indicated. **D.** Percentage of cell death induced by 1 µM staurosporine in binned concentration ranges of the laminin gradient array concentrations (0–1 µg/mL, 1–2 µg/mL, and 2–3.25 µg/mL). Statistical analysis was performed using a Kruskal–Wallis (non-parametric) test for multiple comparisons on unpaired data.

## Discussion

*In vitro* models that can account for the complexity of the ECM have the potential to replicate key pathogenic features observed in specific disease states^9,8^. Such platforms can be used to assess how the microenvironment impacts treatment responses and disease progression. Extrusion bioprinting provides one approach to producing such model systems in an automated fashion, but typically requires the composition of the hydrogel precursor or bioink formulation to be defined prior to initiating printing, resulting in the evaluation of a limited number of formulations^45,46^. This limits the ability of such models to systematically assess how particular hydrogel-incorporated components and related mechanical properties may play a role in specific cell responses. To address this, printing toolheads that permit real-time modification of the hydrogel precursor composition directly prior to extrusion have been developed^30,32–36,47^. In these systems, compositional variation is achieved by dynamically adjusting the relative flow rates of multiple material streams, and active or passive mixing strategies ensure homogenization of the combined streams before extrusion. Studies that apply such systems have established their utility for efficiently combining materials with diverse properties in an automated, efficient and reproducible manner^30,32–36,47^.

Here we describe a programmable bioprinting mixing toolhead, and apply it to produce an *in vitro* array for studying tumor microenvironment effects. The design of this bioprinting mixing toolhead is informed by an active mixing printhead created by Ober et al.^34^. In that design the mixing rod (or impeller) is inserted directly into the extrusion nozzle, while here the mixing chamber is upstream of the extrusion port. Ober et al. provide an in-depth comparative analysis of the mixing dynamics and impact of channel design, flow rates and hydrogel precursor viscoelasticity in the context of active and passive mixing strategies^34^. A consequence of these constraints is that to ensure efficient homogenization of viscous liquids in passive mixers, long mixing channel lengths and geometries designed to increase mixing efficiency are typically required, and mixing will only occur while hydrogel precursors are flowing through the device. In contrast, an active mixing strategy facilitates the decoupling of the injection (and extrusion) flow rates from mixing, and removes some of the design constraints for the geometry of the mixing channel. Consequently, smaller volume chambers can be employed, which may be preferable when aiming to create large arrays of hydrogel combinations for screening purposes. Furthermore, as active mixing of hydrogels will persist in the chamber in the absence of flow, the homogeneity of hydrogel mixtures can be improved by simply prolonging the mixing duration between extrusions, irrespective of flow rates. A related advantage to being able to pause extrusion is that the nozzle can be freely moved to different print locations without risk of hydrogel precursor carry over, which was availed of throughout this study to sequentially print discrete extrusions into the 384 well plate format. Our mixing toolhead also shares characteristics with the recently published two component printhead designed by Teves et al. Their active mixing strategy facilitates on-the-fly adjustments to the composition of multi component materials and they highlight the advantage of being able to tune material compositions between extrusions^36^.

The predicted concentration profiles for hydrogel precursors A and B in the extruded mixtures (Fig. 2) assumes that a homogenous mixture is achieved prior to each new injection. As illustrated in Fig 2, while the length of the mixing rod is matched with the length of the mixing chamber (40 µl), it does not extend into the nozzle (160 µl). Therefore, while the rod ensures active mixing in the mixing chamber, to ensure that a homogenous mixture is formed within the relatively larger volume of the nozzle requires that the vortex created by the rod and the duration between extrusions are sufficient. Mixing in the nozzle will likely also be influenced by laminar flow as the hydrogel precursor mixture that enters the nozzle is expected to undergo axial (Taylor-Aris) dispersion, such that fluid at the center moves faster than fluid near the nozzle walls, and diffusion redistributes molecules radially^48^. The nozzle geometry and shear-thinning properties of hydrogel precursors such as alginate solutions would promote faster central flow and mixing, while with higher viscosity fluids slower diffusion rates would be expected. Therefore, in addition to active mixing in the chamber, passive mixing effects in the nozzle are likely also contributing to the mixing efficiencies observed for the extruded hydrogel precursors.

The concentration profiles observed for the experimental fixed-ratio prints (Fig. 2) largely supported the predicted one-phase exponential increases in the contribution of hydrogel precursor B to the extruded mixtures. These were quantified as a product of the dextran-AF488 fluorescence and counts of fluorescent microspheres. Furthermore, the dextran-AF488 signal and the fluorescent microspheres appeared to be evenly distributed throughout each well, indicating homogenous mixing of hydrogel precursors A and B. These prints were programmed to ensure that after each extrusion, the nozzle remained above the extrusion well for 30 seconds, while mixing continued. This dwell was implemented to ensure sufficient mixing, but also proved practically necessary to ensure that the extruded hydrogel precursor mixture dissociated completely from the nozzle opening, before the nozzle was moved to the next well. However, as noted in the results section, image analysis of the fixed ratio printed arrays revealed that the first detectable contribution of hydrogel precursor B (dextran-AF488 or fluorescent microspheres) to the extruded mixtures appeared typically seven wells later than was predicted by the calculations outlined in Fig. 2. Here we discuss the design and operation features of the mixing tool that may contribute to this lag phase in the extrusion profiles. Firstly, to avoid the inclusion of air bubbles the mixing tool is prefilled with hydrogel precursor A. This includes the inlet channels to which the syringe pump tubing for hydrogel precursors A and B are subsequently connected to (Fig. 2). These inlet channels have a volume of 8.5 µl, and consequently any initial injection of hydrogel precursor B into the toolhead will first displace this prefilled volume of hydrogel precursor A into the mixing chamber, which would account for some delay in the establishment of the expected extrusion profiles. A second consideration is that the prefilling of the system with hydrogel precursor A may introduce a degree of back pressure in the hydrogel precursor B channel that is not immediately overcome by the first hydrogel precursor B injections. As the injection and extrusion channels are open it is reasonable to assume that any such pressure issues would be overcome within the initial injection, but may nonetheless also contribute to the observed lag. Finally, the delay may partially be due to the behavior of hydrogel precursor A present in the nozzle prior to printing and not in direct contact with the active mixing rod. Considering this 160 µl as a tapering column of hydrogel, the upper section will be the first to experience a vortex due to its proximity to the rotating rod. This region of hydrogel precursor A will also be the initial portion into which hydrogel precursor B will be mixed. Therefore, there will be an initial gradient of hydrogel precursor B along the axial length of nozzle, and so the major component of the initial extrusions will be the prefilled hydrogel precursor A in the nozzle. Clearing this entire 160 µl of hydrogel precursor A by plug flow would incur a lag of 8 x 20 µl extrusions. This is more than the experimentally observed lag, suggesting that a degree of hydrogel precursor B does mix with the prefilled hydrogel precursor A, but it is likely a major contributor to the observed delay. Additional discrepancies to the predicted extrusion profile for the alginate-based prints were also observed. For example, steeper concentration gradients than predicted were observed between extruded wells 10 to 20, when the injected volume of hydrogel precursor B was increased (Fig. 2E and F). This may in part be explained by an overestimation of the total mixing volume. As the nozzle threads into the Luer lock and overlaps around the extrusion port of the mixing chamber (Fig. 2A), the available internal volume of the nozzle will be reduced relative to that of an unconnected nozzle. The 0:100 fixed injection ratio prints deviated most from the predicted profiles, and here there is no injections of hydrogel precursor A. It is possible that in the absence of the resistance pressure provided from hydrogel precursor injections through the opposing inlet, there may be a degree of overshoot for hydrogel precursor B injections caused by residual pressure that results in post-injection flow. This would cause more hydrogel precursor B to enter the mixing chamber than that defined by the injection volume. Importantly, while we do not have conclusive explanations for all of the observed discrepancies in the extrusion profiles, the incorporation of fluorescent trackers directly into the hydrogel arrays ensures that the specific contribution of a specific hydrogel precursor can be quantitatively determined and directly correlated with effects observed in any discrete well of the array.

Linear gradients are widely used to screen a continuous range of mixing ratios between two components, and there are microfluidic based solutions that support the formation of gradients with the inclusion of encapsulated cells^32,24,28^. Linear gradient arrays could be generated with the current mixing toolhead by modifying the G-code that controlled the injection ratios, permitting switching between fixed ratio (Fig. 2) and dynamic ratio (Fig. 3) gradient printing programs. The ability to bioprint A_3_I-A mini-spidroin gradients, including A_3_I-A–mCherry and alginate/A_3_I-A combinations (Fig. 4), provides a useful platform for generating hydrogel arrays with tunable biochemical and mechanical properties. This is particularly useful given that recombinant spider-silk hydrogel stiffness is concentration dependent^16,41^, and such arrays could be applied to screen for matrix stiffness-induced effects with relevance to tumor progression and fibrosis. However, the printed A_3_I-A concentration profiles did deviate from the predicted gradients (Fig. 4). The A_3_I-A solutions were more viscous than the alginate and this likely impacted their mixing and dispensing through the toolhead. For example, in the print combining alginate with A_3_I-A-mCherry, the A_3_I-A-mCherry concentration gradient was much steeper than expected, reaching its maximum value by extrusion 50 rather than the expected 90, which resulted in several failed extrusions in the final 35 wells of the array (Fig. 4C). This observed profile suggests that the intended injected volumes of A_3_I-A-mCherry were exceeded, and as discussed above, this may be a result of a residual pressure effect such that A_3_I-A-mCherry continues to flow into the mixing chamber post-injection. Despite this deviation from the expected output, these experiments demonstrate that the mixing toolhead is capable of mixing and extruding highly concentrated protein-solution gradients. Furthermore, the incorporation of fluorescent reporters, as demonstrated here with mCherry, provides a built-in quantitative readout of the actual contribution of a given hydrogel, enabling well-specific definition of the array’s composition. Nevertheless, the fit of these extrusion profiles could likely be improved by adjusting the injection volumes to account for the behavior of high viscosity materials. A similar approach could be extended beyond these mCherry conjugates to generate gradient arrays of A_3_I-A functionalized with other fusion proteins, such as adhesion motifs or enzymes^40^. Furthermore, the capacity to systematically blend fusion and non-fusion forms of A_3_I-A in an automated process may help maintain gelation properties that may be compromised when recombinant spidroins are engineered with fusion proteins^41^.

To demonstrate how bioprinted ECM gradient arrays could be applied to investigate the potential impact of cell-ECM interactions on cell sensitivity to the pro-apoptotic agent staurosporine, we first determined MDA-MB-231 cell death responses to staurosporine gradients in alginate-only hydrogel arrays. Staurosporine induces apoptosis through its ability to inhibit a broad-spectrum of protein kinases^49^. Bioprinting staurosporine gradients directly onto pre-seeded MDA-MB-231 cells proved compatible with cell staining, confocal imaging, and high-content image analysis using CellProfiler. The dose–response curves generated from the array data revealed the EC_50_ for staurosporine-induced MDA-MB-231 cell death in alginate-only arrays to be 1.01 µM, following overnight treatment. This is in line with previous reports from 2D cultures in which staurosporine induced MDA-MB-231 cell death was reportedly at 84% following 48 h exposure to 0.5 µM^50^, and 35% following 24 h exposure to 1 µM^51^. Further, we established that morphological changes indicative of cell rounding were detected at a lower EC_50_ than that for cell death. In future approaches, these bioprinted arrays could be also be subjected to downstream analyses with Cell Painting approaches to study diverse cell responses to libraries of compounds^52^.

Laminin 511 is a constituent of the breast basement membrane, alongside multiple other laminin isoforms. Laminin expression is altered during basement membrane remodeling, which is associated with tumor cell invasion and metastasis, and laminin-511 specifically has been implicated in breast cancer progression^44,53^. Laminin engagement with specific cell surface integrins promotes the recruitment of focal adhesion-associated proteins and activation of downstream signaling pathways that regulate survival, migration, and proliferation^54^. MDA-MB-231 cells revealed increasing sensitivity to staurosporine along the increasing laminin 511 concentration gradient arrays. In contrast, in the absence of staurosporine, MDA-MB-231 cell viability was unaffected. This aligns with previous reports from Vasaturo et al, where a laminin-associated increase in staurosporine-induced death was observed in MDA-MB-231 cells^51^. However, we initially considered that the incorporation of laminin 511 into these tumor cell microenvironments may engage integrin-mediated survival mechanisms and promote resistance to staurosporine-induced death. One possible interpretation is that MDA-MB-231 integrin interactions with laminin 511 selectively increases staurosporine-targeted kinases, thereby increasing cell susceptibility to staurosporine-induced death. Another consideration is the format of the assay, as the laminin 511 gradient is deposited onto MDA-MB-231 cells that are pre-seeded in the 384 well plates and cultured overnight with culture medium. Therefore, these cells have already established attachments and adhesions in part through interactions with ECM molecules present in the culture medium, and produced by the cells themselves. Consequently, the introduction of laminin 511 into the surrounding microenvironment may directly compete with and disrupt pre-existing adhesions and perturb pro-survival signaling mechanisms, in turn rendering cells more sensitive to staurosporine. While future work will be required to elucidate the specific mechanism of laminin-induced sensitivity to staurosporine, these data demonstrate how bioprinted ECM gradient arrays provide a platform for screening the contribution of specific tumor microenvironment components to altered cell sensitivity to anti-cancer treatments.

### Limitations of the current study

To characterize the mixing and extrusion performance of the tool we employed alginate-based hydrogel precursors, which are widely used in various extrusion based bioprinting approaches, but lack cell adhesion motifs that are essential to recapitulate the cell-ECM interactions present in living tissues. However, we established that the mixing tool is not limited to handling alginate hydrogel precursors; for example, by generating arrays with A_3_I-A mini-spidroin protein solutions. Further, while the hydrogel component in our gradient arrays provide a 3D microenvironment through which diffusion of solutes and gases can occur, and ECM molecules can be incorporated, the MDA-MB-231 cell line is cultured under standard 2D conditions, prior to encapsulation under the hydrogel arrays. This facilitates cell imaging, but it makes it difficult to conclusively attribute cell responses to the composition of the gradients alone, as cell attachments in the well will also influence their behavior. Moreover, although CellProfiler enabled high-throughput image analysis, variability in fluorescence intensity between wells, potentially amplified by the presence of the hydrogel, occasionally complicated cell segmentation. This made it challenging to optimize a single set of segmentation parameters applicable across all 90 wells, particularly for elongated cells in control conditions. Furthermore, we limited our studies to the MDA-MB-231 cell type, and additional work with other breast cancer cell lines will be required to determine whether the laminin 511 effects on staurosporine sensitivity are generally applicable or limited to this specific cell type. Finally, the effective range of concentrations that can be screened in a gradient of a drug or ECM molecule of interest is constrained by the maximum concentrations of available formulations and related solubilities from which hydrogel precursor stocks can be prepared. For the laminin 511 gradients presented here, the dilution of laminin-511 in the alginate hydrogel solutions produced a narrow range of screening concentrations in the resulting arrays. Although this limited exploration of a wide concentration range, reports of disease-associated ECM variations suggest that relatively small changes in protein concentration may contribute to relevant changes in cellular behavior and disease progression^55–58^.

In conclusion, we outline the design, fabrication, and operation of an active mixing toolhead for programmable bioprinting of tumor microenvironment arrays. We describe fixed and dynamic ratio printing strategies that facilitate the production of gradient arrays with alginate and A_3_I-A mini-spidroin hydrogels. We provide a quantitative fluorescence-based approach to defining the specific concentrations of hydrogel precursor components in discrete extrusions across the entire gradient, and imaging protocols that are compatible with imaging cell death and morphological responses. Finally, we bioprint a gradient of microenvironments containing increasing laminin-511 concentrations, and determine that the presence of this tumor-associated ECM molecule increases MDA-MB-231 sensitivity to an apoptosis-inducing agent.

## Materials and Methods

### Reagents and chemicals

Sodium alginate (12–40 kDa; Sigma-Aldrich) was used for hydrogel precursor preparation. Alexa Fluor-conjugated dextran (10 kDa; Invitrogen, Thermo Fisher Scientific) was resuspended in phosphate-buffered saline (PBS; Gibco, Thermo Fisher Scientific) to a final concentration of 5 mM and stored at −20 °C. CountBright Plus Absolute Counting Beads (4 µm diameter, 10^5^ beads/50 µL; Invitrogen, Thermo Fisher Scientific) were used to track hydrogel precursor concentrations, and visualize mixing efficiency. Calcium chloride (CaCl_2_, 50 mM in HEPES buffer; Cellink, Sweden) served as the alginate crosslinking solution. Staurosporine (1 mM in DMSO; Abcam) was used to induce apoptosis. Laminin-511 (Biolaminin 511 LN, 0.1 mg/mL in pH 7.2 buffer; BioLamina, Sundbyberg, Sweden) was used to pre-condition alginate hydrogel precursor stocks. For live/dead staining, propidium iodide (1 mg/mL in water; Invitrogen, Thermo Fisher Scientific) was diluted to 2 µg/mL in Opti-MEM prior to use. Calcein-AM Green (1 mg/mL; Invitrogen, Thermo Fisher Scientific) was diluted to 4 µg/mL prior to staining.

### Cell culture

The human breast cancer cell line MDA-MB-231 was obtained from the American Type Culture Collection (ATCC). Cell line authentication was performed by Eurofins Genomics using short tandem repeat (STR) profiling, confirming the identity of the MDA-MB-231 cell line. Cells were cultured in Dulbecco’s Modified Eagle Medium (DMEM) GlutaMAX (Gibco, Thermo Fisher Scientific, Uppsala, Sweden) supplemented with 10% fetal bovine serum (FBS; Thermo Fisher Scientific) and 1% penicillin–streptomycin (PS; Thermo Fisher Scientific). This formulation is referred to hereafter as culture medium. Cells were maintained in a humidified incubator at 37 °C with 5% CO2. For microscopy-based experiments, cells were cultured in phenol red-free Opti-MEM (Thermo Fisher Scientific) supplemented as indicated.

### Alginate preparation

Sodium alginate solutions (5% w/v) were prepared by dissolving alginate powder overnight in either PBS or Opti-MEM at room temperature under continuous magnetic stirring. For assessing hydrogel precursor mixing in the absence of cells, alginate solutions were prepared in PBS. For cell culture experiments, alginate was dissolved in Opti-MEM supplemented with 1% PS under sterile conditions in a laminar flow hood. Following dissolution, the alginate solution was sterile-filtered using a 0.44 µm pore-size filter. Prior to syringe loading, solutions were centrifuged at 1000 × g for 4 min release air bubbles. Additional components, including fluorescent tracking molecules (dextran and microspheres), drugs (staurosporine), or proteins (laminin 511), were incorporated by mixing with the alginate solution using syringes connected via Luer-lock connectors. Following mixing, samples were centrifuged again (1000 × g, 4 min) to remove any newly introduced air bubbles before printing experiments. For cell-based experiments, hydrogel precursor formulations were maintained at 37 °C prior to mounting on the syringe pump system.

### Design, fabrication and operation of the mixing tool

The mixing tool was designed to be used in combination with the open-source E3D tool changer and motion system (E3D-online, London, United Kingdom). This 3D bioprinter has previously been described in detail by Engberg et al.^37^. Computer-aided design (CAD) of the mixing toolhead components was performed using Fusion 360 (Autodesk Inc, San Rafael, USA). All 3D-printed parts for the mixing tool were printed on a Form 3 SLA 3D printer (Formlabs) in standard clear resin (Formlabs) with a Z-resolution of ∼100 μm. Post printing all parts produced were washed and cured according to the manufacturer’s instructions. The flow and volume of hydrogel precursor A and hydrogel precursor B injected from syringes into the mixing chamber were controlled by independent syringe pump tools. The syringe pump tools are controlled by the Duet Wifi controller board and actuated by stepper motors. The 5 mL syringes (Omnifix Luer Lock, VWR) containing hydrogel precursors A or B were connected to their respective inlets on either side of the lower housing of the mixing tool module using fixed lengths of tubing (Masterflex Transfer Tubing, Tygon, VWR) (Fig. 1D). Either end of the respective hydrogel precursor tubing was fitted with a nut and a ferrule (Cole-Parmer). A 3D printed Luer Lock was designed to fit the nut and connect to the syringe pump tool, and the other end connects to the mixing tool inlet via a tubing connector (Fig. 1A). To ensure sealed connections of the nuts to the connecting ports, O-rings (nitrile rubber, 7 x 1.5, inner dia. (mm) x thickness (mm); Dione Kullager AB, Sweden) were fitted in the mixing tool inlets and the syringe connectors. The two inlets connect directly with the mixing chamber, which encloses the metal mixing tool rod. The mixing chamber was sealed at the top using an O-ring (fluoro rubber, 1.43 x 1.5, inner dia. (mm) x thickness (mm); Kullager, Sweden) surrounding the rod. The volume of the sealed mixing chamber is ∼57 mm^3^, and is reduced to ∼34 mm^3^ with the mixing tool rod inserted. The mixing shaft was manufactured either as a single solid piece machined from stainless steel at Uppsala University workshop or as a two-part assembly in which the mixing tool base was redesigned to accommodate a press-fit connection with a 2 mm stainless steel dowel pin (CP 2×32 DIN 7 Stainless Steel Cylindrical Pin, FastPro, Mölndal, Sweden) turned down to 1.5 mm diameter, resulting in a protruding shaft length of 13.5 mm. The extrusion outlet at the bottom of the lower housing of the mixing module was fitted with nozzle for hydrogel precursor extrusion. The mixing tool base was attached to a stepper motor driven by an external single board computer (SBC; Raspberry Pi 5, 8 GB RAM, Raspberry Pi Ltd., Cambridge, UK). The rotation speed (rpm) and rotation time (s) were controlled via a Python script running on the raspberry pi (Supplementary Material and Methods, Fig. S1).

### Quantification of extrusion concentrations for fixed injection ratios

To determine the mixing profile of hydrogel precursor A with hydrogel precursor B for a specific injection ratio, each print was divided into 40 x 20 µl extrusions deposited into 384-well plates (Thermo Scientific, 384-well black plate, optically clear polymer bottom). The G-code for each printed array was generated using a Python script (Supplementary Material and Methods, Fig. S2) with the following input parameters: number of wells desired (40), syringe diameter, volume to be extruded into each well (20 µl), and distance between wells (4.5 mm). The print consisted of a grid of 10 columns by 4 rows, with a dwell time of 30 s between each extrusion. The E3D tool changer was programmed to follow a serpentine path moving along the row direction. The dwell time allowed the viscous hydrogel precursor mixture to be fully extruded from the nozzle prior to the tool head moving to the next well. Hydrogel precursor A corresponded to a 5% (w/v) sodium alginate solution in PBS and hydrogel precursor B corresponded to a 5% (w/v) sodium alginate solution in PBS conditioned with 2 μM dextran Alexa Fluor 488 and 14% (v/v) CountBright plus absolute counting beads. The following static ratios of hydrogel precursor A: hydrogel precursor B (Hy. A: Hy. B) were printed: 70:30, 50:50, 30:70, and 0:100. The mixing tool was primed with hydrogel precursor A prior to printing. When consecutive prints with different fixed injection ratios were performed, the mixing chamber was purged and refilled with hydrogel precursor A between each print. Extrusion concentrations in each well were quantified by image analysis, using dextran Alexa Fluor 488 and fluorescent microbeads as proxies for the hydrogel precursor B content in each mixture. The fluorescent microbeads were counted using the ImageJ (Fiji)^59^ particle analysis plugin, while the concentration of dextran Alexa Fluor 488 was determined by measuring fluorescence intensity from images acquired from each well, using an LSM700 confocal microscope (Zeiss, Jena, Germany) equipped with a 5X objective (Zeiss) to visualize the entire well area. Microbeads and dextran were imaged over three optical sections using the 555 nm and 488 nm lasers, respectively. The experimental data were compared with the extrusion profiles predicted for mixtures of hydrogel precursor A and hydrogel precursor B generated with the different injection ratios (as outlined in Fig. 2B).

### Quantification of extrusion concentrations for dynamic injection ratios

The dynamic injection ratio printing protocol was designed to generate a near linear extrusion profile enabling the fabrication of a gradient array covering hydrogel B concentrations from 0-100%. These arrays were divided into 90 wells with 20 µl of mixed hydrogel precursor extruded per well, and as described above, the mixing chamber is primed with hydrogel precursor A. The G-code for these dynamic gradient prints encoded the Hy. A: Hy. B injection ratios and the number of extruded wells per ratio, as follows: 85:15 (12 wells), 80:20 (4 wells), 75:25 (4 wells), 70:30 (4 wells), 65:35 (4 wells), 60:40 (4 wells), 55:45 (4 wells), 50:50 (4 wells), 45:55 (4 wells), 40:60 (4 wells), 35:65 (4 wells), 30:70 (4 wells), 25:75 (4 wells), 20:80 (4 wells), 15:85 (8 wells), and 0:100 (18 wells). The dynamic gradient print is composed of 16 different fixed Hy. A: Hy. B injection ratios. Confocal microscopy imaging of dextran-AF488 intensity and fluorescent microspheres number was performed as described above for fixed mixing ratio. Plate reader imaging of dextran-AF 488 intensity was performed on a BioTek Synergy H4 Hybrid Reader using the Gen5™ software. Data points corresponding to failed extrusion events, where no alginate solution was delivered to the wells, were excluded from the analysis.

### Dynamic gradient printing of A_3_I-A mini-spidroin hydrogel precursor arrays

The A_3_I-A mini-spidroin protein and the mCherry fusion A_3_I-A mini-spidroin (A_3_I-A-mCherry) protein was recombinantly expressed and purified as previously reported^40,41^. The dynamic injection ratio print procedure with 16 fixed injection ratios (described above) was followed to extrude gradient arrays of A_3_I-A mini-spidroin solutions in a 384 well plate format. One exception applies for the A_3_I-A (200 mg/ml): A_3_I-A 200 (mg/ml) + dextran (Fig. 4A), which was performed with 20 fixed ratios, as follow: 95:5 (4 wells), 90:10 (4 wells), 85:15 (4 wells), 80:20 (4 wells), 75:25 (4 wells), 70:30 (4 wells), 65:35 (4 wells), 60:40 (4 wells), 55:45 (4 wells), 50:50 (4 wells), 45:55 (4 wells), 40:60 (4 wells), 35:65 (4 wells), 30:70 (4 wells), 25:75 (4 wells), 20:80 (4 wells), 15:85 (4 wells), 10:90 (4 wells), 5:95 (4 wells) and 0:100 (14 wells). Prior to printing, wells were filled with 20 µL of filtered 20 mM Tris HCl buffer (pH 8). For all prints, the syringe containing A_3_I-A mini-spidroin solutions was placed in an ice bath to maintain low temperature during printing to prevent gelation. To condition the viscous A_3_I-A solution with dextran Alexa Fluor 555 (dextran-AF55) the spidroin stock was divided evenly between two syringes. Dextran-AF555 was collected into one syringe, both syringes were connected with a Luer lock, and a homogenous mixture with a final dextran-AF555 concentration of 5 µM was produced by passing the solutions back and forth between the syringes on ice. The dextran-AF555 concentration in this stock was 5 µM and the A_3_I-A concentration was 200 mg/ml. The syringe was centrifugated at 900 rpm for 4 minutes to remove air bubbles. The A_3_I-A and A_3_I-A-mCherry stock concentrations used in the respective dynamic gradient prints are indicated in Fig. 4. The dynamic gradient prints with A_3_I-A solutions were imaged with the plate reader or by confocal microscopy using 5X or 10X objectives, and z-stacks containing 3 planes were collected per well. The z-stacks were converted to maximum intensity projections in ImageJ and the average fluorescence per well was quantified. As for the alginate prints, control prints were prepared directly from the individual syringes containing the stock solutions to quantify the maximum fluorescence for dextran Alexa Fluor 555 conditioned A_3_I-A solutions, or of the mCherry fusion in the A_3_I-A-mCherry.

### Printing gradient hydrogel precursor arrays of increasing staurosporine concentrations on MDA-MB-231 cells

MDA-MB-231 cells were seeded in 384 well plate at a density of 45,000 cells/cm^2^ which corresponds to 3150 cells/well, and cultured in culture medium under standard conditions (5% CO_2_, 37 °C) for one day prior to printing. Prior to printing, the culture medium in the 384 well plate was exchanged for 20 µl of Opti-MEM supplemented with 5% FBS and 1% PS. Alginate was prepared in Opti-MEM supplemented with 1% PS. Staurosporine stocks were prepared in DMSO (1mM), and to avoid DMSO-induced crosslinking of the alginate^60^, a 142.9 µM solution of staurosporine was first prepared in Opti-MEM prior to mixing with the alginate solution. The hydrogel precursor B alginate solution was mixed with 14% (v/v) of 142.9 µM staurosporine solution and 1% dextran Alexa Fluor 680 (10 kDA) using two syringes connected with a Luer Lock adaptor. The hydrogel B stock contained 20 µM staurosporine. The hydrogel precursor A alginate solution was mixed with 14% (v/v) of DMSO % in Opti-MEM to control for the presence of these reagents in hydrogel precursor B. The dynamic gradient print protocol was performed as outlined above, and printing was performed at room temperature. The extrusion in well 90 corresponds to the last position of the mixing tool, where residual hydrogel flow in the system may lead to occasional over-extrusion, resulting in volumes exceeding 20 µL and staurosporine concentrations above the studied range. Data from overfilled well 90 were excluded from analyses of staurosporine concentration, cell death response, and aspect ratio analysis.

### Printing gradient hydrogel precursor arrays of increasing laminin 511 concentrations on MDA-MB-231 cells

Cell seeding and handling was performed as described above in the staurosporine printed gradient array section. Alginate was prepared in Opti-MEM supplemented with 1% PS, and the hydrogel precursor B preparation was mixed with 14% (v/v) of 0.1 mg/ml laminin 511 solution (10% glycerol, 2% sodium azide) and 1% dextran Alexa Fluor 680 (10 kDA) using two syringes connected with a Luer Lock adaptor. The laminin concentration in the hydrogel precursor B was 14 µg/ml. The alginate solution for hydrogel precursor A was mixed with 14% (v/v) of glycerol 10% in PBS to account for the presence of this reagent in hydrogel precursor B. After printing, alginate crosslinking and washing step, a solution of 1 µM staurosporine or a control treatment was added to each well (50 µl).

### Cell viability controls

To monitor the effect of the printing conditions alone on cell viability (i.e. independent of hydrogel precursor extrusion); cell seeded wells were included in the 384 well array plate and maintained without hydrogels precursor in either 20 µl OptiMEM (5% FBS, 1%PS) or 50 µl of culture medium, throughout the duration of the printing process.

### Single syringe pump extrusions for hydrogel precursor A and hydrogel precursor B controls

Following gradient array printing with the mixing tool, the hydrogel precursors A- and B-containing syringes were disconnected from the mixing tool and directly fitted with a nozzle. Ten wells (20 µl/well) of each hydrogel precursor were printed in the 384 well plate. These control wells were used to determine the average maximum bead number and dextran fluorescence for hydrogel B, from which the hydrogel B percentage in each well of the gradient array are determined. For staurosporine and laminin in alginate gradient prints, these controls enabled the determination of hydrogel B concentrations in each well of the gradient array using dextran AF-680 fluorescence intensity. Additionally, these control extrusions, printed on pre-seeded MDA-MB-231permit cell viability evaluation below hydrogel A and B to be assessed independently of the mixing process.

### Hydrogel precursor cross-linking post printing

After the 90 well dynamic injection ratio prints (45 min), the 384 well plate was first incubated at 37 °C for 10 min. Subsequently, each well was incubated with 30 µl of 50 mM calcium chloride (CaCl_2_) for 2 min 30 sec to cross-link the alginate hydrogel precursor. Following cross-linking, 40 µl of Opti-MEM (5% FBS, 1 %PS) was added to each well to dilute the CaCl_2_ and was then aspirated from each well. Opti-MEM (5% FBS, 1% PS) was added to each well (50 µl) to culture cells overnight.

### Staurosporine and laminin concentration calculations

For the staurosporine and laminin gradient prints (Fig. 5 and Fig. 6), the average autofluorescence of hydrogel A control wells (without dextran) was subtracted from both the fluorescence values of the gradient prints and the average fluorescence of hydrogel B control wells (with dextran) prior to concentration calculations. Negative values resulting from this background correction were set to zero. The maximum staurosporine and laminin concentrations were defined using the hydrogel B control wells. As 20 µL of hydrogel precursor containing either 20 µM staurosporine or 14 µg/mL laminin was printed into wells containing a total volume of 90 µL, complete diffusion throughout the well corresponded to maximal concentrations of 4.44 µM staurosporine and 3.11 µg/mL laminin. In some cases, calculated concentrations within the gradient exceeded these theoretical maxima due to variability and averaging of the fluorescence measurements obtained from the hydrogel B control wells.

### Cell staining, imaging, and analysis

The Opti-MEM (5% FBS, 1% PS) solution on top of the hydrogel was removed without disrupting the gels. A staining solution of propidium iodide and calcein in Opti-MEM was added to each well (50 µl) and incubated for 3h. The staining solution was not washed before imaging. Each well was imaged using a 20X objective on a LSM700 (Zeiss) with the tile-scan function. Tile scans were reconstituted as one image for each well as an hyperstack and all generated hyperstack are concatenated in a stack using Image J. The three channels were split: propidium iodide (nucleus of dead cells), calcein (cytoplasm of live cells), dextran AF 680 (proxy for hydrogel B concentration). An Image J macro was used to label each well and save it as an independent tif format image using the following labelling: ““Plate1_” + well_number + “_site0_Ch1.tif“”, for each channel, modifying Ch1 to CH_2_ and CH_3_ when appropriate. Image analysis was performed using CellProfiler (v4.2.8, www.cellprofiler.org, Broad Institute, Cambridge, MA, USA), an open-source high throughput image analysis software^61^. All images were then imported in CellProfiler and metadata of images were extracted from the file names using the following expression: ^(?P<PLATE>.*)_(?P<WELL>W[0-9](*1*))_(?P<SITE>site[0-9]+)_(?P<CHANNEL>Ch[0-9]+)\.tif. A pipeline was built to identify the live cells with the Calcein AM channel (IdentifyPrimaryObject) and identify the dead cells with the propidium iodide channel (IdentifyPrimaryObjects). The IdentifyPrimaryObject function permits cells to be segmented from the images and quantify cell counts for each channel per well. Based on the live cell objects identified, a mask image of the dextran channels without the live cells was created to avoid calcein stained cells contribution in the dextran channel, using the MaskImage module. A measure image intensity module (MeasureImageIntensity) was used to measure dextran fluorescence intensity in each masked image. Shape descriptors for live and dead cells are measured using the MeasureObjectSizeShape. The aspect ratio was calculated through CalculateMath module using the AreaShape category of shape descriptors by dividing the MajorAxisLenght by the MinorAxisLength. Several iterations in the test mode were run to find optimal module parameters identifying live and dead cells. The results were exported as a cvs format file. For control prints consisting of laminin only, additional image-processing steps were incorporated into the pipeline to improve segmentation of live cells. In the absence of staurosporine, the low levels of cell death, higher cell density, and elongated cell morphology made segmentation more challenging, particularly when combined with uneven illumination within tile-scan images and variations in fluorescence intensity between wells. Illumination correction was performed within each well using the CorrectIlluminationCalculate and CorrectIlluminationApply modules. The IdentifySecondaryObjects module was added to refine the cytoplasmic segmentation of live cells, which are initially detected as primary objects from the calcein channel. This step improved segmentation robustness under conditions of variable fluorescence intensity between wells by extending object boundaries to include lower-intensity cytoplasmic regions.

### Statistical analysis

Statistical analyses were performed using Graphpad Prism 10 software. The specific tests applied and the number of replicates included are indicated in the associated figure legends.

## Data availability

All data supporting the findings of this study are available on reasonable request to the corresponding author.

## Acknowledgements

The authors acknowledge funding provided by the European Union’s Horizon 2020 Research and Innovation Programme to the project ‘BioTriB’, grant agreement No. 956004. This study was conducted as part of the Additive Manufacturing for the Life Sciences (AM4Life) consortium, which is funded by Sweden’s Innovation Agency Vinnova (grant number 2019-00029), and by additional grants from the Swedish Cancer Society (Cancerfonden; grant numbers 20 1285 PjF and 23 2692 Pj 01 H), and the Göran Gustafsson’s Foundation (Göran Gustafssons Stiftelser). This work was also supported by FORMAS (2023-00871) to B.S; and Olle Engkvists Stiftelse (233-0334), Knut and Alice Wallenberg Foundation (2023.0331), FORMAS (2023-01313), and the Swedish Research Council (2024-02919) to AR. 3D printing was performed at U-PRINT: Uppsala University’s 3D-printing facility at the Disciplinary Domain of Medicine and Pharmacy and SciLifeLab Uppsala. We would also like to thank Palash Chandravanshi and Malin Marcström for their contributions testing earlier prototypes, as well as Olle Pontén for writing the Python script controlling the stepper motor rotation.

## Author contributions

M.M. Conceptualization, methodology, investigation, formal analysis, visualization, writing – original draft, writing – review and editing; E.S. Investigation, validation, writing – review and editing; A.E. Conceptualization, methodology, investigation, writing – review and editing; C.S. Methodology, Writing – review and editing; F.H. Methodology, investigation, writing – review and editing; T.B.P. Resources, methodology, writing – review and editing; B.S. Resources, methodology, writing – review and editing; A.R. Resources, supervision, writing – review and editing; J.K. Conceptualization, resources, supervision, funding acquisition, writing – review and editing; P.O.C. Conceptualization, validation, visualization, resources, supervision, writing – original draft, writing – review and editing. All authors approved the final draft for submission.

## Declaration of AI use

The authors acknowledge the use of generative artificial intelligence (AI) tools, including OpenAI ChatGPT (GPT-5.5 release) and Microsoft Copilot for language editing and manuscript refinement. All scientific content, interpretations, analyses, and conclusions in the final manuscript are the authors own.

