## Supplementary information for "Programmable bioprinting of tumor microenvironment arrays reveals laminin-dependent drug sensitivity"

Supplementary Material and Methods

**Python script for controlling mixing rotor**

Custom Python script running on a Raspberry Pi developed to control a stepper motor (NEMA 11) through a driver (DRV8825). Adjustable parameters include the rotational speed (RPM), mixing duration, and start delay. The script converts the selected RPM into motor step delays based on the configured microstepping mode and executes motor movement through the RpiMotorLib library. Additional functions include testing the motor speed, stopping the motor rotation and optional smooth transitions between rotational speeds during operation.

```
from tkinter import *
from tkinter import ttk
import time
import RPi.GPIO as GPIO
# import the library
from RpiMotorLib import RpiMotorLib

global MotorRunning,changeRPM, currentRPM, currentStepDelay, runDuration, stop
MotorRunning = False
changeRPM = False
currentRPM = 100
currentStepDelay=0
runDuration=60
stop = False

#define GPIO pins
GPIO_pins = (18, 15, 14) # Microstep Resolution MS1-MS3 -> GPIO Pin
direction= 21       # Direction -> GPIO Pin
step = 20      # Step -> GPIO Pin

global StepSize, UpdateFrequency, SPP
StepSize = "1/4"
UpdateFrequency = 1
SPP = {"Full" : 200, "Half" : 400, "1/4" : 800, "1/8" : 1600, "1/16":3200}
# Declare an named instance of class pass GPIO pins numbers
Nema11_Mixer = RpiMotorLib.A4988Nema(direction, step, GPIO_pins, "DRV8825")

def go_at_RPM(nRPM,duration=60):
    global MotorRunning, RPM, currentRPM, runDuration, root, resetTimer, stop
    runDuration = duration
    RPM = nRPM
    run=True
    verbose = False
    steps = int(float(SPP[StepSize])/UpdateFrequency)
    stepDelay = 1/(nRPM/60*SPP[StepSize]*2.2)
    #Lazy solution
    timestamp = int(time.time())
    while(run and not stop):
        MotorRunning = True
        Nema11_Mixer.motor_go(False, StepSize, steps,  stepDelay, False, stepDelay)
        root.update()
        resetTimer.set(f"Remaining time: {int((runDuration+timestamp)-time.time())}")
        #Keep going until interrupted if runduration negative
        if(int(time.time())>runDuration+timestamp and runDuration > 0):
            run = False
            MotorRunning = False
        if(changeRPM):
            RPM = currentRPM
            stepDelay = 1/(nRPM/60*SPP[StepSize])
    MotorRunning = False

def change_RPM(newRPM, newDuration, changeDelay):
    timestamp = int(time.time())
    globals()['stop'] = False
    if(not globals()["MotorRunning"]):
        while(timestamp+changeDelay>int(time.time())):
              time.sleep(0.1)
              globals()["root"].update()
        globals()['stop'] = False
        go_at_RPM(newRPM,newDuration)
    elif(globals()["smoothChange"]):
        globals()["changeRPM"] = True
        globals()["currentRPM"] = newRPM
        globals()["runDuration"] = newDuration
    else:
        if(int(time()) > timestamp+changeDelay):
            globals()["currentRPM"] = newRPM

def stop_stepper():
    globals()['stop'] = True

def test_speed(RPM,duration = 15):
    global MotorRunning, root
    run=True
    #4000 steps = 100 rounds
    steps = int(float(SPP[StepSize])/UpdateFrequency)
    stepDelay = 1/((RPM*SPP[StepSize])/60)
    timestamp = int(time.time())
    while(run):

        MotorRunning = True
        Nema11_Mixer.motor_go(False, StepSize , steps,  stepDelay, False, stepDelay)

        if(int(time.time())>duration+timestamp):
            run = False
    MotorRunning = False

def driver_cleanup():
    GPIO.cleanup()

if __name__ == '__main__':
    global root, resetTimer
    GPIO.setmode(GPIO.BCM)
    root = Tk()
    root.title("RPI: U-PRINT mixing tool")
    pad_x = 5
    pad_y = 10
    content = ttk.Frame(root)
    RPM_ent = ttk.Entry(content, width = 8)
    runtime_ent = ttk.Entry(content, width = 8)
    delay_ent = ttk.Entry(content, width=8)
    curRPM = StringVar(root,"Current RPM: 0")
    resetTimer = StringVar(root,"Remaining time: 0")
    setRPM_lbl = ttk.Label(content,text="Set RPM:")
    setRuntime_lbl = ttk.Label(content,text="Set Runtime:")
    setDelay_lbl = ttk.Label(content,text="Set Start Delay:")
    curRPM_lbl = ttk.Label(content, textvariable = curRPM)
    resetTimer_lbl = ttk.Label(content, textvariable = resetTimer)


    global smoothChange
    smoothChange = BooleanVar(value=True)
    smoothchanges_check = ttk.Checkbutton(content, text="Smooth RPM change", variable=smoothChange)

    Test_btn = ttk.Button(content, text="Test", command = lambda: test_speed(int(RPM_ent.get())))
    Stop_btn = ttk.Button(content, text="Stop", command = lambda: stop_stepper())
    Run_btn = ttk.Button(content, text="Run", command = lambda: change_RPM(int(RPM_ent.get()),int(runtime_ent.get()),int(delay_ent.get())))


    content.grid(column=0, row=0,ipadx = pad_x,ipady = pad_y)
    curRPM_lbl.grid(column=0,row=0,padx = pad_x,pady=pad_y )
    resetTimer_lbl.grid(column=0,row=1,padx = pad_x,pady=pad_y)
    smoothchanges_check.grid(column=0,row=2, padx=pad_x,pady=pad_y)
    setRPM_lbl.grid(column=1, row=0,padx = pad_x,pady=pad_y)
    setRuntime_lbl.grid(column=1, row=1,padx = pad_x,pady=pad_y)
    setDelay_lbl.grid(column=1, row=2,padx = pad_x,pady=pad_y)
    RPM_ent.grid(column=2, row=0,padx = pad_x,pady=pad_y)
    runtime_ent.grid(column=2, row=1,padx = pad_x,pady=pad_y)
    delay_ent.grid(column=2, row=2,padx = pad_x,pady=pad_y)

    Test_btn.grid(column=3, row=0,padx = pad_x,pady=pad_y)
    Stop_btn.grid(column=3, row=1,padx = pad_x,pady=pad_y)
    Run_btn.grid(column=3, row=2,padx = pad_x,pady=pad_y)
    content.pack(padx = pad_x,pady = pad_y)

    root.resizable(False,False)
    root.protocol("WM_DELETE_WINDOW",driver_cleanup())
    root.mainloop()
    Nema11_Mixer.motor_stop()
```

**Graphical user interface for G-code generation**

Custom Python script providing a graphical user interface (GUI; Supplementary Fig. 1) for generating G-code to control the extrusion of defined volumes of hydrogel precursor into user-specified wells within a multi-well plate. Adjustable parameters include syringe type, extrusion volume per well (μL), pause time (s), well range, and positioning coordinates. The script calculates the required extrusion distance based on syringe dimensions and generates the corresponding G-code for automated dispensing.

```
import tkinter as tk
from tkinter import messagebox
from tkinter import filedialog
from math import pi
import pyperclip

def well_to_coords(well, well_plate_size, start_x, start_y):
    row = ord(well[0]) - ord('A')
    col = int(well[1:]) - 1
    x = start_x - row * well_plate_size
    y = start_y - col * well_plate_size
    return (x, y)

class Application(tk.Frame):
    def __init__(self, master=None):
        super().__init__(master)
        self.master = master
        self.grid()
        self.create_widgets()

    def create_widgets(self):
        self.lbl_syringe_id = tk.Label(self, text="Syringe ID")
        self.lbl_syringe_id.grid(row=0, column=0)
        self.syringe_var = tk.StringVar(self)
        self.syringe_var.set("250uL")
        self.opt_syringe_id = tk.OptionMenu(self, self.syringe_var, "250uL - Hamilton", "1000uL -  Hamilton", "5mL - Fischerbrand", "5mL - Omnifix", "3mL - Omnifix")
        self.opt_syringe_id.grid(row=0, column=1)

        self.lbl_volume = tk.Label(self, text="Extruded Volume per Well (µl)")
        self.lbl_volume.grid(row=1, column=0)
        self.txt_volume = tk.Entry(self)
        self.txt_volume.grid(row=1, column=1)

        self.lbl_pause = tk.Label(self, text="Pause (s)")
        self.lbl_pause.grid(row=2, column=0)
        self.txt_pause = tk.Entry(self)
        self.txt_pause.grid(row=2, column=1)

        self.lbl_start_well = tk.Label(self, text="Start Well")
        self.lbl_start_well.grid(row=3, column=0)
        self.txt_start_well = tk.Entry(self)
        self.txt_start_well.grid(row=3, column=1)

        self.lbl_end_well = tk.Label(self, text="End Well")
        self.lbl_end_well.grid(row=4, column=0)
        self.txt_end_well = tk.Entry(self)
        self.txt_end_well.grid(row=4, column=1)

        self.lbl_wells_per_row = tk.Label(self, text="Wells per Row")
        self.lbl_wells_per_row.grid(row=5, column=0)
        self.txt_wells_per_row = tk.Entry(self)
        self.txt_wells_per_row.grid(row=5, column=1)

        self.lbl_z_movement = tk.Label(self, text="Z Movement (mm)")
        self.lbl_z_movement.grid(row=6, column=0)
        self.txt_z_movement = tk.Entry(self)
        self.txt_z_movement.insert(0, "12")
        self.txt_z_movement.grid(row=6, column=1)

        self.lbl_plate_size = tk.Label(self, text="Centre to centre well (mm)")
        self.lbl_plate_size.grid(row=7, column=0)
        self.txt_plate_size = tk.Entry(self)
        self.txt_plate_size.insert(0, "4.5")
        self.txt_plate_size.grid(row=7, column=1)

        self.lbl_start_x = tk.Label(self, text="Start Coordinate X")
        self.lbl_start_x.grid(row=8, column=0)
        self.txt_start_x = tk.Entry(self)
        self.txt_start_x.insert(0, "293.1")
        self.txt_start_x.grid(row=8, column=1)

        self.lbl_start_y = tk.Label(self, text="Start Coordinate Y")
        self.lbl_start_y.grid(row=9, column=0)
        self.txt_start_y = tk.Entry(self)
        self.txt_start_y.insert(0, "139.4")
        self.txt_start_y.grid(row=9, column=1)

        self.btn_generate = tk.Button(self)
        self.btn_generate["text"] = "Generate GCode"
        self.btn_generate["command"] = self.generate_gcode
        self.btn_generate.grid(row=10, column=0, columnspan=2)

        self.btn_copy = tk.Button(self)
        self.btn_copy["text"] = "Copy to Clipboard"
        self.btn_copy["command"] = self.copy_to_clipboard
        self.btn_copy.grid(row=11, column=0, columnspan=2)

        self.btn_save = tk.Button(self)
        self.btn_save["text"] = "Save as GCODE"
        self.btn_save["command"] = self.save_as_gcode
        self.btn_save.grid(row=12, column=0, columnspan=2)

        self.txt_gcode = tk.Text(self)
        self.txt_gcode.grid(row=13, column=0, columnspan=2)

    def generate_gcode(self):
        try:
            # Calculate E value
            syringe_id = self.syringe_var.get()
            if syringe_id == "250uL - Hamilton":
                syringe_diameter = 2.3  # Corrected diameter for 250uL
            elif syringe_id == "1000uL - Hamilton":
                syringe_diameter = 4.61
            elif syringe_id == "5mL - Omnifix":
                syringe_diameter = 12.35
            elif syringe_id == "3mL - Omnifix":
                syringe_diameter = 9.45
            else:  # 5mL - Fischerbrand
                syringe_diameter = 11.73


            volume = float(self.txt_volume.get())  # µl
            pause = int(float(self.txt_pause.get()) * 1000)  # ms

            e_value = volume / (pi * (syringe_diameter / 2) ** 2)
            well_plate_size = float(self.txt_plate_size.get())
            start_x = float(self.txt_start_x.get())
            start_y = float(self.txt_start_y.get())

            # Get wells
            start_well = self.txt_start_well.get().upper()
            end_well = self.txt_end_well.get().upper()

            # Get number of wells per row
            wells_per_row = int(self.txt_wells_per_row.get())

            # Get Z Movement
            z_movement = self.txt_z_movement.get()

            # Generate gcode
            gcode = []
            current_row = start_well[0]
            current_col = int(start_well[1:])

            direction = 1  # forward

            while current_row <= end_well[0]:
                for _ in range(wells_per_row):
                    current_well = current_row + str(current_col)
                    x, y = well_to_coords(current_well, well_plate_size, start_x, start_y)
                    gcode.append(f"G1 X{x} Y{y} F1000")  # Move to the well
                    gcode.append(f"G1 Z-{z_movement} F1000")  # Move down
                    gcode.append(f"G92 E0")  # Set current extruder position to 0
                    gcode.append(f"G1 E{e_value} F100")  # Extrude
                    gcode.append(f"G4 P{pause}")  # Pause
                    gcode.append(f"G1 Z{z_movement} F1000")  # Move up

                    if current_well == end_well:
                        break

                    # Only change column if we are not at the end of the row
                    if _ != wells_per_row - 1:
                        current_col += direction

                if current_well == end_well:
                    break

                current_row = chr(ord(current_row) + 1)  # Move to the next row

                direction *= -1  # Change direction

            # Show gcode
            self.txt_gcode.delete(1.0, tk.END)
            self.txt_gcode.insert(tk.END, "\n".join(gcode))
        except Exception as e:
            messagebox.showerror("Error", str(e))

    def copy_to_clipboard(self):
        pyperclip.copy(self.txt_gcode.get(1.0, tk.END))

    def save_as_gcode(self):
        filename = filedialog.asksaveasfilename(defaultextension=".gcode")
        with open(filename, 'w') as f:
            f.write(self.txt_gcode.get(1.0, tk.END))

root = tk.Tk()
app = Application(master=root)
app.mainloop()
```

Supplementary Figure S1.

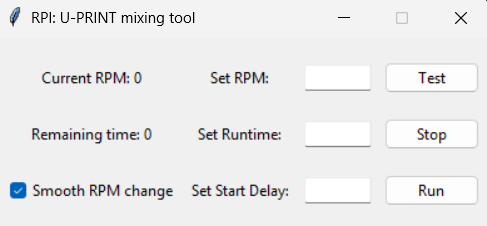

**Fig. S1.** Graphical user interface of the Python script used to control stepper motor operation on the Raspberry Pi. The interface allows the user to define the rotational speed (RPM), runtime (s), and start delay (s), and includes options for smooth RPM change, as well as controls for test, stop and run motor rotation, with real-time display of current RPM (default: 0) and remaining time (default: 0).

Supplementary Figure S2.

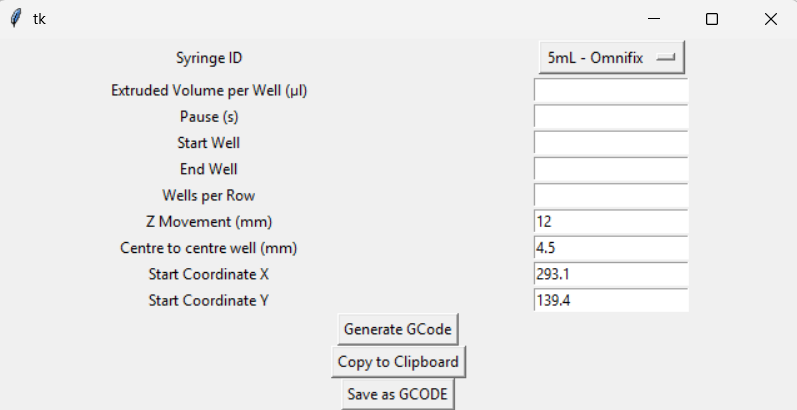

**Supplementary Figure 2.** Graphical user interface of the Python script used to generate G-code for extrusion into multi-well plates. The interface allows the user to define the syringe type (default: 5 mL Omnifix), extrusion volume per well, pause time (s), start and end wells, number of wells per row, Z-axis displacement (default: 12 mm), center-to-center well spacing (default: 4.5 mm for a 384-well plate), and starting X and Y coordinates (default: X 293.1, Y 139.4).
